# A connectomic resource for neural cataloguing and circuit dissection of the larval zebrafish brain

**DOI:** 10.1101/2025.06.10.658982

**Authors:** Mariela D. Petkova, Michał Januszewski, Tim Blakely, Kristian J. Herrera, Gregor F.P. Schuhknecht, Robert Tiller, Jinhan Choi, Richard L. Schalek, Jonathan Boulanger-Weill, Adi Peleg, Yuelong Wu, Shuohong Wang, Jakob Troidl, Sumit Kumar Vohra, Donglai Wei, Zudi Lin, Armin Bahl, Juan Carlos Tapia, Nirmala Iyer, Zachary T. Miller, Kathryn B. Hebert, Elisa C. Pavarino, Milo Taylor, Zixuan Deng, Moritz Stingl, Dana Hockling, Alina Hebling, Ruohong C. Wang, Lauren L. Zhang, Sam Dvorak, Zainab Faik, Kareem I. King, Pallavi Goel, Julian Wagner-Carena, David Aley, Selimzhan Chalyshkan, Dominick Contreas, Xiong Li, Akila V. Muthukumar, Marina S. Vernaglia, Teodoro Tapia Carrasco, Sofia Melnychuck, TingTing Yan, Ananya Dalal, James M. DiMartino, Sam Brown, Nana Safo-Mensa, Ethan Greenberg, Michael Cook, Samantha Finley-May, Miriam A. Flynn, Gary Patrick Hopkins, Julie Kovalyak, Meghan Leonard, Alanna Lohff, Christopher Ordish, Ashley L. Scott, Satoko Takemura, Claire Walsh, John J. Walsh, Daniel R. Berger, Hanspeter Pfister, Stuart Berg, Christopher Knecht, Geoffrey W. Meissner, Wyatt Korff, Misha B. Ahrens, Viren Jain, Jeff W. Lichtman, Florian Engert

**Affiliations:** Department of Molecular and Cellular Biology, Center for Brain Science, Harvard University, Cambridge, MA 02138, USA; Google Research, Zürich 8002, Switzerland; Google Research, Boulder, CO 80301, USA; Sorbonne Université, CNRS, Inserm, Institut de la Vision, F-75012 Paris, France; Department of Visual and Data-Centric Computing, Zuse Institute Berlin (ZIB), Berlin, Germany; School of Engineering and Applied Sciences, Harvard University, Cambridge, MA 02138, USA; Department of Biology, University of Konstanz, Konstanz, Germany; Harvard College, Harvard University, Cambridge, MA 02138, USA; Harvard Graduate School of Design, Harvard University, Cambridge, MA 02138, USA; Brandeis University, Waltham, MA 02454, USA; University of Massachusetts Dartmouth, Dartmouth, MA 02747, USA; Google Research, Mountain View, CA 94043, USA; Janelia Research Campus, Howard Hughes Medical Institute, Ashburn, VA 20147, USA; Harvard Department of Stem Cell and Regenerative Biology, Harvard University, Cambridge, MA 02138, USA

**Author notes:** These authors contributed equally to this work (Mariela D. Petkova, Michał Januszewski).

## Abstract

We present a correlated light and electron microscopy (CLEM) dataset from a 7-day-old larval zebrafish, integrating confocal imaging of genetically labeled excitatory (*vglut2a*) and inhibitory (*gad1b*) neurons with nanometer-resolution serial section EM. The dataset spans the brain and anterior spinal cord, capturing >180,000 segmented soma, >40,000 molecularly annotated neurons, and 30 million synapses, most of which were classified as excitatory, inhibitory, or modulatory. To characterize the directional flow of activity across the brain, we leverage the synaptic and cell body annotations to compute region-wise input and output drive indices at single cell resolution. We illustrate the dataset’s utility by dissecting and validating circuits in three distinct systems: water flow direction encoding in the lateral line, recurrent excitation and contralateral inhibition in a hindbrain motion integrator, and functionally relevant targeted long-range projections from a tegmental excitatory nucleus, demonstrating that this resource enables rigorous hypothesis testing as well as exploratory-driven circuit analysis. The dataset is integrated into an open-access platform optimized to facilitate community reconstruction and discovery efforts throughout the larval zebrafish brain.

## Introduction

Brain function emerges from dense networks of synaptically connected neurons. Synaptic connectivity likely serves as the fundamental substrate for information processing and behavioral control. While this principle has long been appreciated, only recently has it become feasible to map synaptic connections at scale across brain regions ^1–4^ and, in some cases, entire brains ^5–9^. Among available techniques, volumetric electron microscopy (vEM) is the benchmark method capable of resolving all synaptic contacts, tracing fine-caliber neurites through the dense neuropil, and revealing the intracellular specializations and organelles that differentiate cell types. However, brain-wide vEM connectomes are currently only available in the nematode and the fruit fly, where EM based circuit analysis is starting to be leveraged at scale for biological discovery ^10–15^. Still, these volumes often lack *in situ* molecular annotations and omit the peripheral organs that receive sensory input or carry out motor output, limiting their interpretability.

We address these limitations in the larval zebrafish (*Danio rerio*), currently the only vertebrate model of sufficiently small size to allow for the application of brain-wide vEM and to permit tracing of synaptic pathways from peripheral sensory and autonomic organs into and throughout the brain. The study of these connectivity patterns enables structural analysis of how inputs are transformed into motor and visceral outputs. All brains are characterized by recurrent networks in which feedback loops are ubiquitous and critical for efficient control ^12,16–20^, and where inhibitory signals often play a decisive role in regulating activity ^21,22^. Determining not just the presence of connections but also their polarity - excitatory or inhibitory - is therefore essential for understanding how activity propagates through the brain. Here we have leveraged the larval zebrafish’s small size and optical transparency, which have made it a workhorse for circuit neuroscience and brain-wide functional imaging ^23,24^, to provide a synaptic resolution anatomical mapping of excitatory and inhibitory circuits across the entire brain.

To facilitate and democratize these efforts we present and share a multimodal, brain-wide dataset from an intact 7-day-old larval zebrafish that combines vEM with confocal imaging of genetically labeled glutamatergic and GABAergic neurons. The dataset includes automated segmentation, molecular annotation of over 40,000 neurons, and 30 million synapses between axons and dendrites with inferred polarity for 21 million of these. It covers the brain and anterior spinal cord, capturing circuitry from peripheral input through central processing to motor and autonomic output.

We demonstrate the utility of this resource through three circuit vignettes focused on sensory processing, decision making, and internal state regulation. In the *posterior lateral line system*, we trace how directionally tuned hair cells convey water flow information to hindbrain neurons, revealing an analogue representation with a chiral bias to counterclockwise flow. In the *anterior hindbrain*, we reconstruct motifs supporting motion integration, including recurrent excitation and contralateral inhibition consistent with models of evidence accumulation. In the *tegmentum*, we map long-range projections from an excitatory cluster which is analogous to the mammalian periaqueductal grey, to refute and refine existing models of its role in regulating arousal, motor state, and visceral responses to threat. Beyond these examples, the dataset is integrated into an online platform <u>(website</u>) that is optimized for community-driven circuit reconstruction and discovery, enabling new insight into the structural logic of vertebrate brain function.

## Results

### Correlated light and electron microscopy reveals neurotransmitter identity across the zebrafish brain

To relate neuronal neurotransmitter subtypes to ultrastructural anatomy across an entire brain, we combined confocal light microscopy (LM) and serial section electron microscopy (ssEM) in the same 7-day post-fertilization (7dpf) zebrafish (**Fig.1A-B**). Excitatory and inhibitory neurons were labeled by transgenic differential fluorescent expression of *vglut2a* and *gad1b* reporters, respectively, and imaged, in live animals, using high-resolution confocal microscopy. This fish was then processed for ssEM, producing a nearly complete brain volume at 4×4×30 nm resolution (**Fig.1C**, top left). This 370 TB EM dataset captured fine-scale ultrastructural features such as synaptic connections and subcellular structures.

**Fig. 1.**
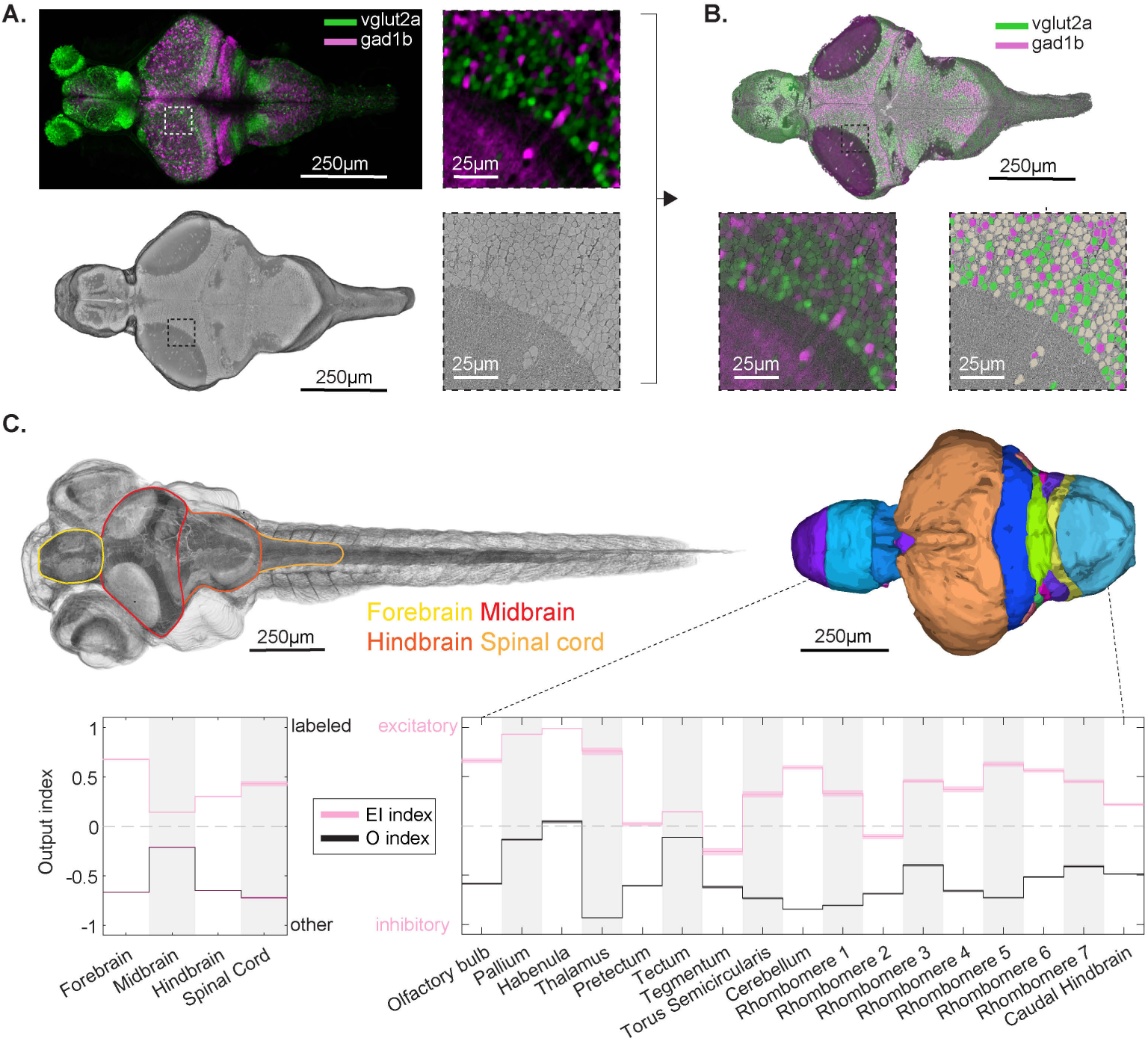
Whole-brain correlated light and electron microscopy (CLEM) enables mapping of molecularly defined neuronal classes and their anatomical distributions in zebrafish. A, Maximum intensity projection of a confocal light microscopy (LM) volume from a 7 dpf zebrafish brain, showing *vglut2a*-expressing (excitatory, green) and *gad1b*-expressing (inhibitory, magenta) neurons. The same specimen was subsequently imaged with serial section electron microscopy (EM), digitally sectioned to reveal internal structures. Side panels show corresponding LM and EM regions. B, Registered and overlaid LM and EM volumes. Bottom left panel shows the raw overlay of fluorescent LM signal on EM ultrastructure; the bottom right panel shows the resulting cell-type annotations: *vglut2a* (green), *gad1b* (magenta), and unlabeled (tan). C, Rendering of the entire fish EM volume annotated with the major anatomical boundaries (left) and segmented into finer subdivisions (right). Cell-level output drive per region is summarized by two indices: *excitatory-inhibitory* index (EI, pink) reflecting relative polarity among labeled neurons, and *other* index (O, black) reflecting the overall balance of labeled versus other cells. Shaded regions in the Manhattan plots indicate bootstrapped standard error of the mean (SEM).

Registering the LM and EM datasets involved three key challenges. First, alignment was complicated by the relative sparsity of labeled cells in LM compared to the complete dense cell representation in EM. Second, the cytoplasmic nature of fluorescence in LM made it difficult to resolve individual neurons in densely packed regions. Third, spatial distortions introduced during fixation, resin embedding, heavy metal staining, tissue sectioning, image stitching and volume alignment led to non-rigid deformations, even though both datasets originated from the same brain. To overcome these obstacles, we developed a multi-step co-registration pipeline that integrated manual annotation, automated alignment, and expert validation.

To align cell identities across the light and electron microscopy volumes, we first performed a coarse registration using anatomical fiduciaries visible in both modalities, such as prominent vasculature and uniquely shaped neurons. We then addressed the multimodal challenge by segmenting both datasets into point clouds of cell locations that could be directly compared to each other (see Methods). Confocal data proved difficult to segment due to dense packing and diffuse cytoplasmic fluorescence, so we adopted a conservative approach that minimized false positives, yielding a point cloud of 35,748 LM-labeled neurons. In contrast, EM segmentation was aided by high contrast and uniformly labeled membranes, allowing us to identify 187,053 cell bodies distributed across the brain, spinal cord, and peripheral ganglia. Of these, we estimated 116,000 to be neurons - a lower bound (see next section)-with the remainder likely to be glia or immature neurons.

We refined the alignment through iterative point-cloud matching between the EM and LM centroids, establishing 9,421 unambiguous correspondence points (“landmarks”) across the brain (**SFig.1-2**). Within a 10µm radius of each landmark, EM and LM segmentations could be matched and directly compared. To further assess match quality and assign neurotransmitter identity, we manually reviewed a total of 70,000 (20µm x 20µm) LM/EM overlay images centered on EM cells near landmarks, identifying those with detectable *vglut2a* or *gad1b* expression (**SFig.3-4**). This process yielded 41,175 EM-resolved and fluorescently labeled neurons: 26,915 *vglut2a-*positive and 14,510 *gad1b*-positive cells. Finally, we estimated that the brain contains ∼58,000 *vglut2a-* and *gad1b-*expressing neurons, where ∼70% of these are matched to the EM volume (see Methods). The remaining ∼30% lie outside of the 10 µm matching radius, in regions too distant from any landmark for reliable alignment.

To quantify cell-type composition across brain regions, we introduced two indices, each characterizing the frequency of relative neurotransmitter cell types in each region. First, we defined an overall labeling index (O), which reflects the fraction of neurons that are labeled by canonical excitatory (*vglut2a*) and inhibitory (*gad1b*) markers versus the populations of “other” cell types such as glycinergic, cholinergic, or neuromodulatory neurons and glia, which remained invisible in our transgenic approach. Second, the excitatory-inhibitory index (EI) reflects the normalized difference between the 41,000 validated *vglut2a* and *gad1b* neurons, and it serves to quantify the E/I balance across brain regions, which were identified by registering the vEM volume into the Z-brain atlas ^25^.

Consistent with previous gene expression and anatomical datasets ^25–28^, we observed excitatory dominance in most regions including the pallium, optic tectum and habenula (**Fig. 1C**, bottom). In contrast the tegmentum exhibited a marked inhibitory bias, aligning with known clusters of GABAergic neurons in this ventral midbrain region. Together, the O and EI indices provide a comprehensive, spatially resolved view of neuronal identity that both validates prior anatomical expectations and reveals region-specific gaps in molecular coverage.

### Automated neurite segmentation and brain-wide synapse annotation

To enable detailed EM segmentation and reconstructions of neurons, glia and other cell types we employed a combination of automated and manual methods, tailored to the varying tissue types and tissue preservation quality across different regions (**Fig. 2A**). To that end, the brain and spinal cord were first automatically segmented using a flood-filling network (FFN) pipeline ^29^, generating supervoxels with a negligible rate of falsely merged objects in the brain. In the spinal cord, the quality of tissue preservation progressively declined along the direction of the tail, leading to a decrease in the segmentation quality caudally. Supervoxels throughout the volume were automatically agglomerated into neurite fragments. A total of 180,249 of these fragments contained a soma (**Fig. 2B**), and added up to 51m of neurite length, which was primarily composed of dendrites and represented approximately one-fifth of the total neurite path length in the brain (**SFig. 5**). Compared to previously released zebrafish datasets ^9,30^, our automated segmentation demonstrated improved performance (**SFig. 5**). Common image artifacts and segmentation errors impacting the automated results are documented in **SFig. 5** and further described in the Methods section. This automatically segmented dataset is still fragmented and does not include any fully reconstructed neurons.

**Fig. 2.**
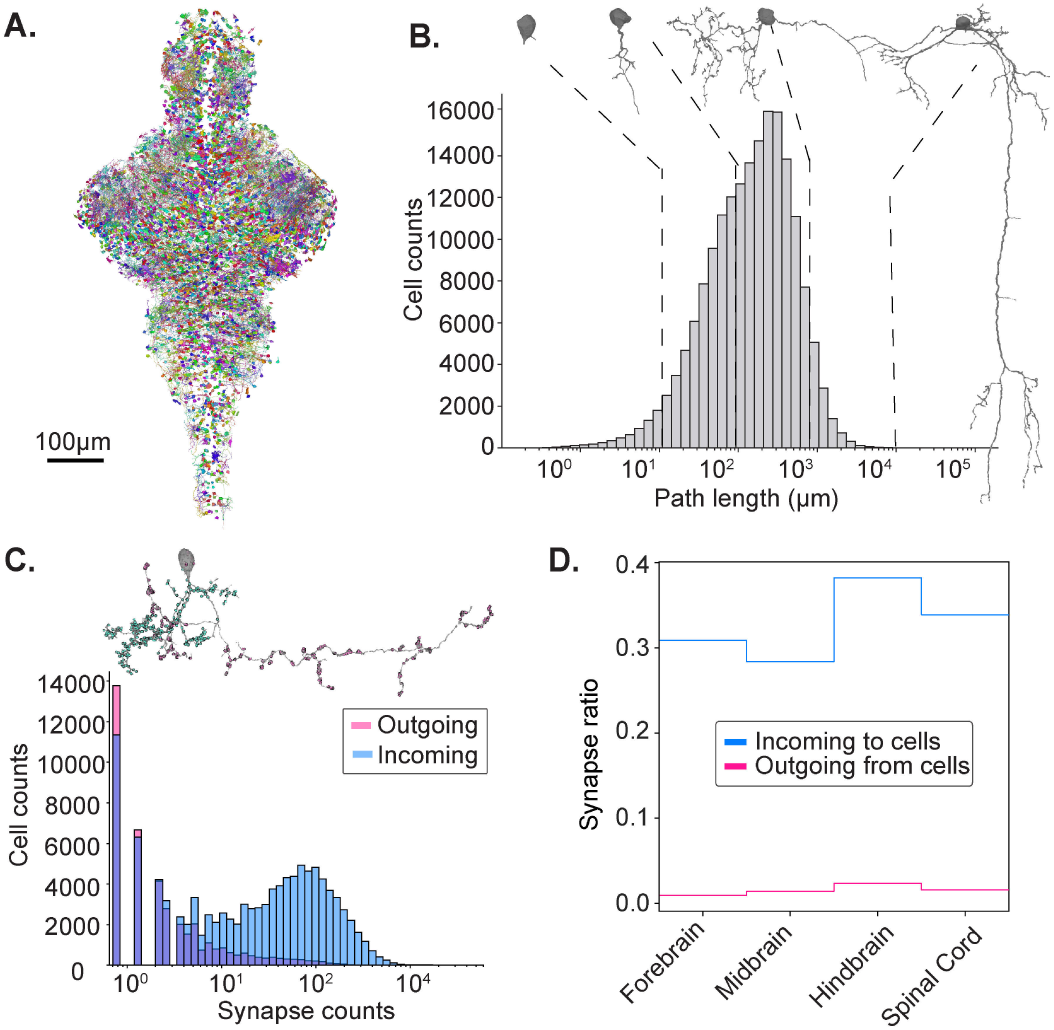
Morphological and synaptic properties of automatically segmented neurons across the brain volume. A, Rendering of a randomly selected 2% of all automatically segmented cells in the volume. **B,** Histogram of neurite path lengths for all segmented cells with representative morphologies shown for key path length regimes. **C,** *Top:* Example neuron showing predicted incoming (pink) and outgoing (blue) synapses. *Bottom*: Log-scale histogram of synapse counts across somata-associated segments; *n* = 108,805 and *n* = 41,930 soma are associated with incoming and outgoing synapses, respectively. **D,** Fraction of synapses captured by somas across the major brain regions, showing a strong bias toward incoming synapses on dendrites.

**Fig. 3.**
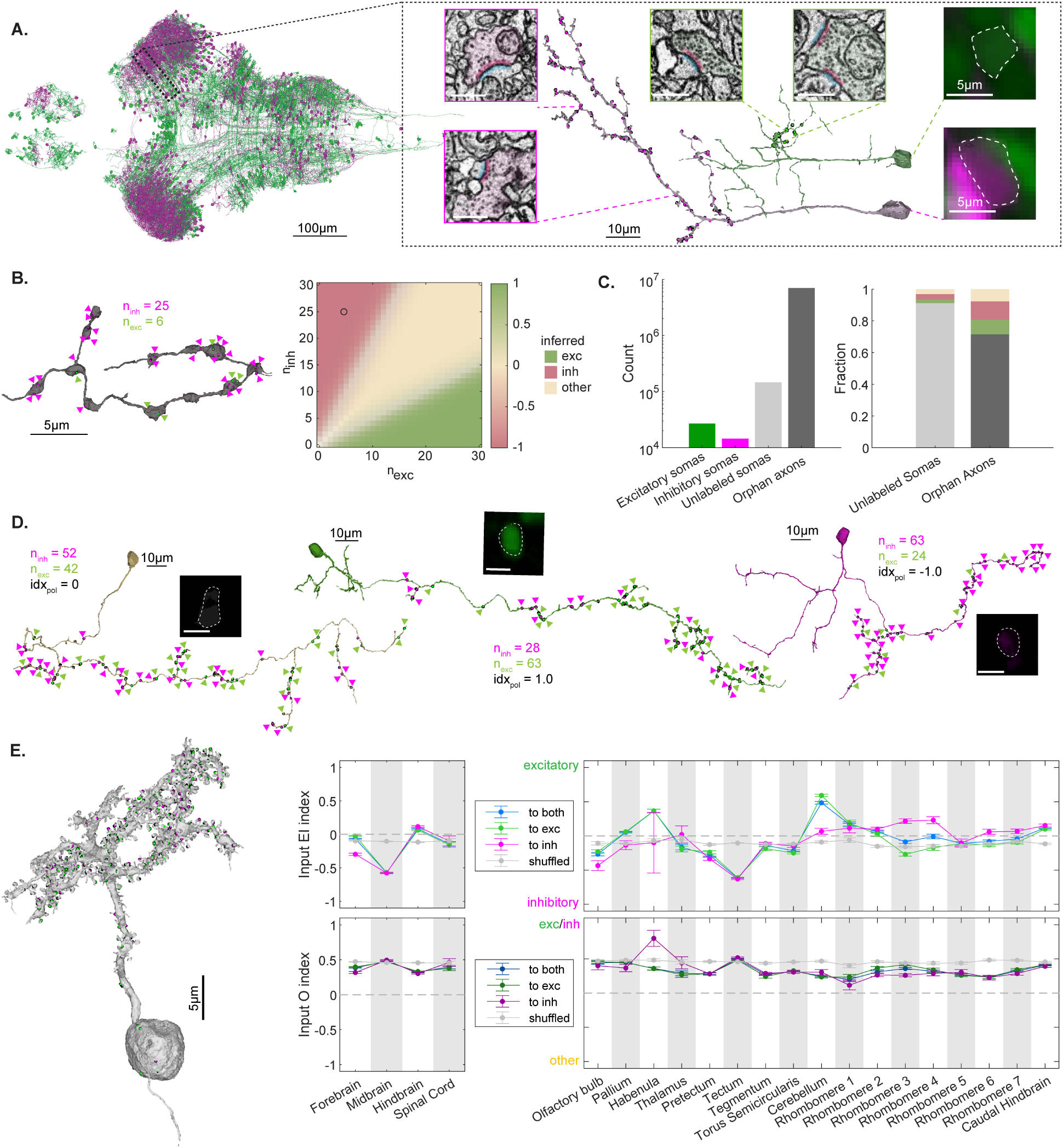
Synapse polarity and distribution across the brain volume. A, Rendering of all neurons used as ground truth to train a synaptic classifier for *vglut2a* (green) and *gad1b* (magenta) synapses with two example neurons selected on the right. Insets show synapses from each of the neurons (scale bar 500nm) as well as the LM overlay at their soma locations (dashed outlines). **B,** Left: Example axon fragment with raw synapse predictions as excitatory and inhibitory (colored triangles). Right: The grid plot shows a certainty-modulated polarity index, *idx_pol_* = *P_exc_* − *P_inh_*, for synapse-type classification based on the number of outgoing excitatory (x-axis) and inhibitory (y-axis) synapses. Hue encodes the direction and magnitude of the bias, while saturation reflects the certainty of the assignment (maximum of *P_exc_, P_inh_,P_other_*). The black circle marks the synaptic configuration of the example axon, which is classified as inhibitory. This visualization supports axon relabeling based on Dale’s rule. **C,** Left: counts of the labeled portions of the dataset. Right: Relabeled distribution of previously unlabeled somas and orphan axons after applying Dale’s rule on segments with at least 4 synapses. **D,** Three examples of manually reconstructed axons for soma that belong to serotonergic, glutamatergic and GABAergic neurons, respectively. The LM overlays, total synapse counts and their polarity index are shown. The colored triangles at synapse locations are sparsified to accommodate the zoom factor. **E,** Left: Incoming raw synapse assignments onto the spiny dendrites of a cell in the cerebellum (green and magenta circles). Right: incoming drive index to 41,000 validated GABAergic and glutamatergic neurons segregated by coarser (left side) and fine-grained (right side) regions in the brain. The top row shows the balance between excitatory and inhibitory input; the bottom row shows the balance between fast-acting (exc/inh) input and modulatory (other) input. Error bars are SEM.

To facilitate computer-assisted manual segmentation and circuit analysis of the automatically segmented data, we provide a collaborative proofreading platform CAVE ^31^ which allows for the manual breaking of rare incorrect links, or adding new ones, while the underlying supervoxels remain fixed (see Methods). We find that this approach enables the reconstruction of cells at an average rate of 10 min to 1 hr per cell, estimated by proofreading ∼1500 cells for specific circuit investigations as showcased in **Figures 4, 5** and **6** below.

**Fig. 4.**
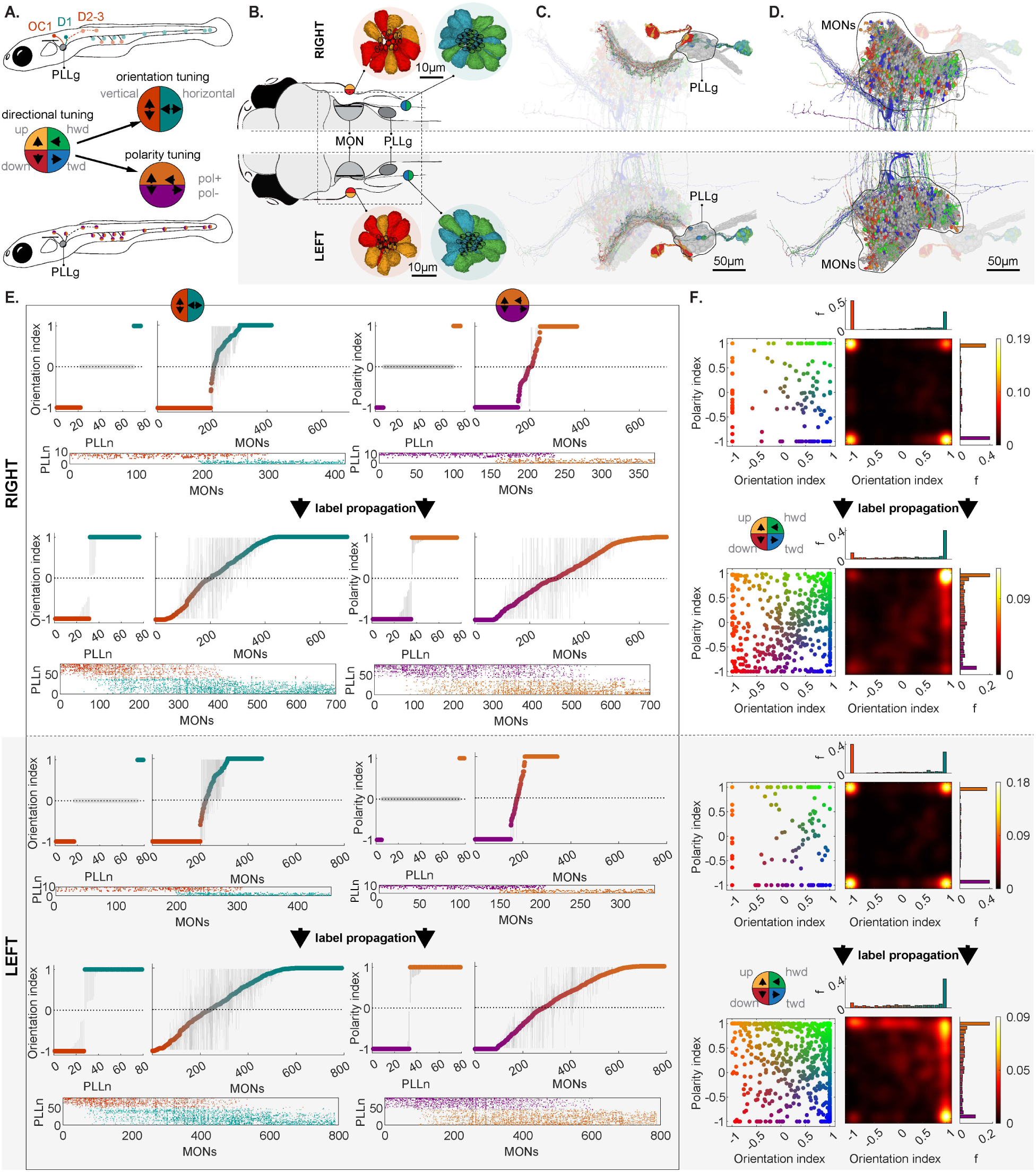
Feedforward transformation of labeled-line directional tuning into a combinatorial flow code in the hindbrain. A, Schematic of directional tuning in the posterior lateral line (PLL) system of larval zebrafish. Neuromasts are selectively tuned to either vertical (vermillion) or horizontal (teal) flow depending on their body position (top), and each neuromast contains mirror-symmetric hair cells that discriminate positive (pol⁺: headward/upward, brown) versus negative (pol⁻: tailward/downward, purple) flow within an axis (bottom). Together, orientation and polarity determine directional tuning. **B,** Hair cells (colored by directional tuning) in two exemplar neuromasts on each side of the fish. Each directional tuning vector corresponds to a specific orientation–polarity pair. **C,** Lateral line afferents (PLLn neurons) receive input from hair cells of a single tuning type and project to the medial octavolateralis nucleus (MON) in the hindbrain. Example neurons are tuned to all cardinal directions (hwd/twd/up/down in green, blue, orange, red). **D,** Automatic reconstruction of MON neurons shows axons of PLLn neurons (colored by tuning) and dendritic arbors of MONs (or colored by connectivity, gray are not connected to the known PLLn). **E.** Bootstrap-based label propagation framework for assigning orientation and polarity tuning to partially labeled PLLn and MON neurons. Top: Initial orientation indices, polarity indices and connectivity matrix for MONs computed from a small, labeled subset of PLLns. Bottom: Final inferred labels for the full population after propagation. Unshaded sets of panels show data from the right side of the fish; gray shaded panels show data from the left side of the fish. The error bar for each neuron represents classification uncertainty, defined as 1−P, where P is the fraction of bootstrap runs assigning that neuron to a given group. **F.** Joint distribution of orientation and polarity tuning in MON neurons. Left: Each MON is plotted by its computed indices, colored by inferred direction preference. Right: Corresponding heat maps show the density of MONs in tuning space, revealing a triangular distribution with hemisphere-specific asymmetries. Right-side MONs show enrichment in tailward directions, while left-side MONs emphasize headward tuning - consistent with net sensitivity to counterclockwise flow around the body.

**Fig. 5.**
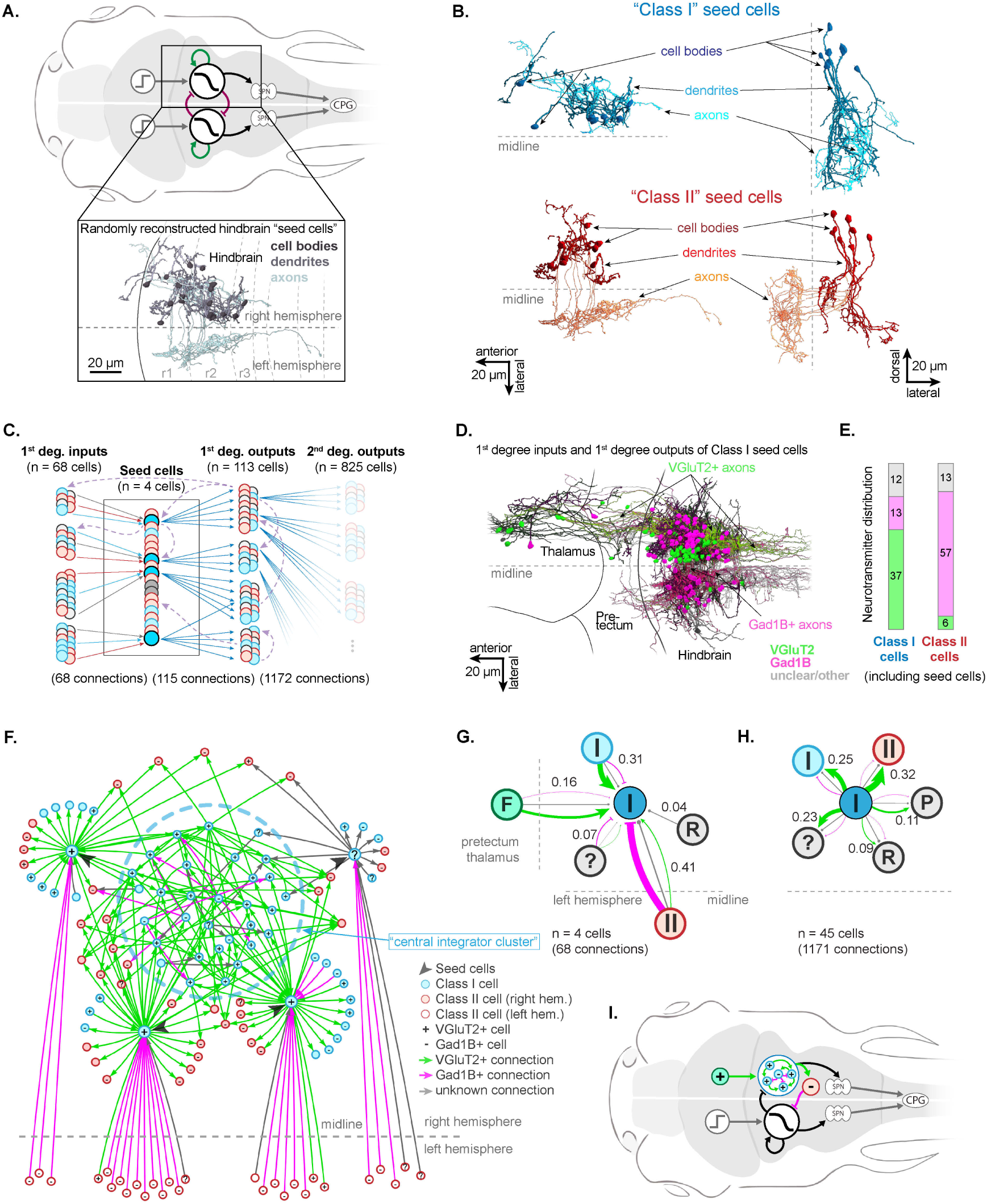
Connectomic analysis of hindbrain integrator circuit. A, Top, hypothetical circuit model for evidence integration, consisting of a putatively recurrent circuit that low-pass filters noisy direction-selective visual evidence from the tectum and pretectum (green arrows). Swim-direction is determined via an inhibitory inter-hemispheric push-pull mechanism (red arrows; “SPN”, spinal projection neurons; “CPG”, central pattern generators). Bottom, top-down projection of 20 randomly reconstructed “seed cells” in the right hindbrain, rhombomeres (r) are indicated. **B,** Seed cells could be divided anatomically into an ipsilateral-projecting (“Class I”; top) and contralateral projecting class (“Class II”, bottom); left panels, top-down projections; right panels, coronal projections from the back. **C,** Schematic depiction of our strategy for exhaustive circuit reconstruction of inputs and outputs of four Class I seed cells (blue, Class I cells; red, Class II cells; grey, other cell classes; solid arrows, reconstructed synaptic connections (same color code); dashed arrows, recurrent ‘within-network’ connections revealed by circuit reconstructions. **D,** Top-down projection showing 1st degree inputs and outputs of the four Class I seed cells; cells are colored by their neurotransmitter identity. **E,** Distribution of neurotransmitter identity for Class I and Class II cells across 1st degree inputs and and 1st degree outputs. **F,** Network graph showing the local synaptic circuit that Class I cells are embedded in. Note that seed cells are connected with each other via a highly recurrent central integrator cluster. Connections are color-coded by neurotransmitter type. **G,** Quantitative input map for four Class I seed cells. **H,** Quantitative output map for Class I cells (seed cells and 1st degree outputs). **G-H,** “I”, Class I cell; “II”, Class II cell; “F”, frontal projection neurons, “R”, raphe neuron; “P”, posterior-projecting cell classes, “?” other cell classes; fractions of all synaptic inputs indicated; connections are colored by neurotransmitter type, line thicknesses indicate fraction of synapses. **I,** Circuit model in **A** updated with tested and confirmed anatomy of the anterior hindbrain (colored in, top).

**Fig. 6.**
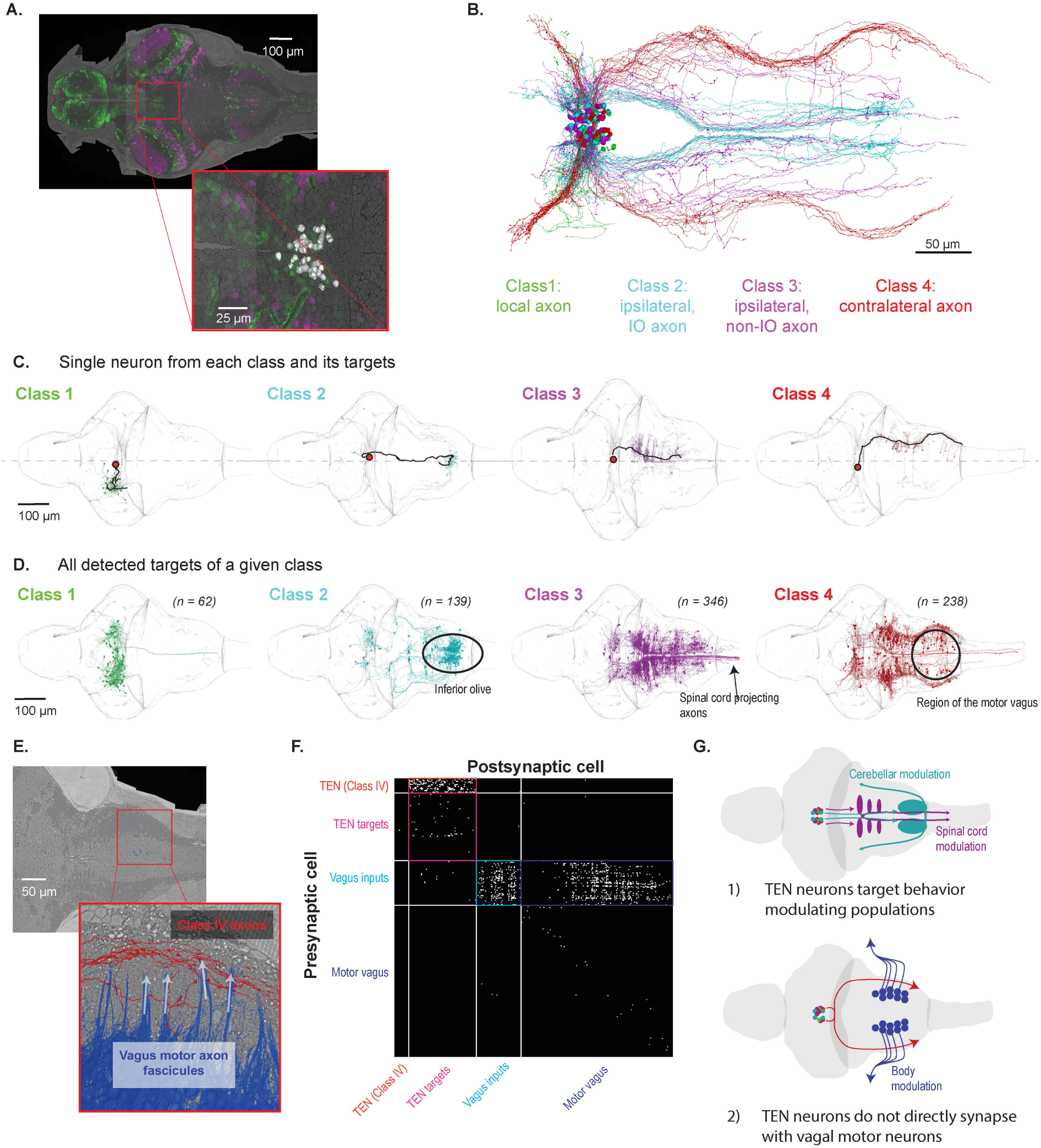
Reconstruction of descending outputs of a tegmental excitatory nucleus (TEN) A, Identification of excitatory nucleus within the midbrain based on excitatory fiduciary. Inset includes overlay of reconstructed cell-bodies (white). **B,** Reconstruction of 111 neurons from the TEN colored by morphological class, **C,** Example neuron from each trace (black), with cell body and axon highlighted, and criteria-meeting postsynaptic neurons from that neuron, **D,** All post-synaptic neurons from each class from synapses filtered by volume and part of a soma-containing segment. **E,** Location of type IV axons (red) relative to the outgoing vagus motor nerve (blue). **F,** Connectivity matrix showing connections among class IV TEN neurons (red), target neurons of class IV TEN neurons (magenta), inputs to motor vagal cells (cyan), and motor vagal neurons (blue). Connectivity is seen in the colored squares indicating TEN to targets (red square), TEN targets to other TEN targets (magenta), vagal inputs to other vagal inputs (cyan), and from vagal inputs to vagal motor neurons (blue). **G,** Summary of findings from TEN reconstructions. Cartoons depicting discovered innervation of behavioral modulating regions (top) and lack of connections between TEN and the motor vagus (bottom).

The non-neuronal body of the zebrafish was not segmented and rare falsely merged supervoxels in the brain and spinal cord might require manual corrections. Both tasks can be performed using VAST ^32^, which supports precise voxel-level annotation for reconstruction at the tissue, cellular or ultrastructural level. While edits made in CAVE are globally visible to the entire community, manual edits in VAST remain private to the user who performs them. This capability ensures that even challenging regions can be accurately segmented, complementing the automated methods used for other parts of the dataset.

We next incorporated automated synapse annotations across the entire dataset. We identified a total of 29.5M axon-to-dendrite and 9,5M axon-to-axon synaptic connections based on pre-and postsynaptic compartment identity derived from semantic masks (see Methods). We estimate a false positive rate of approximately 15%, based on manual review. We next attempted to match, where possible, the presynaptic/postsynaptic terminals to the cell bodies where the axon/dendrite originated. Since a much larger fraction of dendrites than axons are traced to their corresponding cell bodies by automated segmentation, we find a substantially higher ‘incoming’ synaptic count than outgoing counts in our data (**Fig. 2C**). **Figure 2D** summarizes the fraction of synapses assigned to somatic dendrites and axons across the major brain regions. On average, over 30% of all synapses in a region are attributed to dendritic compartments of soma-associated cells, whereas fewer than 2% are attributed to soma-associated axons.

Finally, we can use the presence or absence of synapses on all soma-associated objects to classify all of the 187,000 cells into neurons and glia. We find that 110,000 soma-associated objects receive at least one *synaptic input* onto compartments classified as dendritic by our semantic masks, while an additional 6,000 soma associated objects produce at least one *synaptic output* from compartments classified as an axonal (but otherwise have no incoming synapses). We classify these 116,000 soma-associated objects as neurons, which is a lower bound for the neuron number, because the automatic segmentation does not attach axonal and/or dendritic compartments to every soma. The remaining 71,000 cell count is therefore an upper bound for the number of glia, as it is expected to contain such poorly segmented cell bodies and other non-glial cell types.

### Neurotransmitter inference reveals regional synaptic input patterns in brain-wide EM

We next classified synapses as excitatory, inhibitory or modulatory, enabling quantification of their respective contributions across brain subregions. In mammalian brains, synaptic ultrastructure correlates with neurotransmitter type: glutamatergic synapses typically exhibit asymmetric postsynaptic densities, whereas GABAergic synapses show symmetric densities ^2,33,34^. In contrast, zebrafish synapses do not follow such clear morphological rules. A similar challenge arose in the *Drosophila* connectome, where convolutional neural networks successfully classified synapses into six neurotransmitter types, demonstrating the feasibility of structure-based prediction from EM images ^35^.

To overcome the lack of established morphological correlates of synaptic function in zebrafish, we took advantage of the 41,000 validated *vglut2a*-and *gad1b*-expressing neurons distributed across all brain regions. Of these, 491 *vglut2a-* and 547 *gad1b-*positive neurons were inspected for the presence of an axon with no merge errors, yielding a ground truth set of over 69,000 synapses (31,013 from *vglut2a* and 38,312 from *gad1b*) (**Fig. 3A**). We used this labeled dataset to train a synapse-level convolutional network classifier based on ultrastructural features, which achieved 83% accuracy on held-out data.

To apply this classifier to the full set of ∼30 million synapses between axons and dendrites, it is essential to account for a third class of synapses that are neither excitatory nor inhibitory. A naïve, wholesale assignment of excitatory or inhibitory labels to all remaining synapses would misrepresent this “other”, likely modulatory neurotransmitter synaptic category. Instead, we leveraged Dale’s principle, which posits that all synapses from a given axon segment should share the same primary neurotransmitter identity. In the LM dataset, we observed co-expression of *vglut2a* and *gad1b* exceedingly rarely, suggesting that a dual-transmitter identity ^36^ is not a prominent feature in this system. Under this assumption, an axon composed predominantly of synapses classified as excitatory (or inhibitory) should itself be excitatory (or inhibitory). Although we cannot definitively confirm neurotransmitter identity for axons with a balanced mix of predicted excitatory and inhibitory synapses, the classifier’s 83% synapse-level accuracy suggests that a true excitatory or inhibitory axon should exhibit a clear skew in polarity. Axons with a near-even mix are therefore unlikely to result from misclassification alone and are better interpreted as belonging to an “other” category - encompassing primarily modulatory axons or those with too few synapses to support a confident assignment. We formalized this logic using a Bayesian inference framework (Methods), which estimates the posterior probability of each axon belonging to one of the three classes based on the composition of its predicted synapses. An example inference for an axon segment is shown in **Fig. 3B**.

Using this approach, we extended the classification to all unlabeled parts of the dataset by aggregating synapse-level predictions at the level of axon fragments and somata. In total, the dataset includes approximately 140,000 unlabeled somas and over 7 million orphan axon fragments with at least one synapse. We performed the three-class inference for all axons and somas with ≥4 associated synapses, a threshold at which classification accuracy is meaningful: for example, *P*(exc|*n*_exc_=4)=0.87, P(inh|*n*ᵢₙₕ=4) = 0.87, *P*(other|*n*_exc_=2, *n*ᵢₙₕ=2) = 0.55. This yielded new excitatory, inhibitory, or “other” assignments for 15,000 somas and 2.6 million axon fragments. Although only ∼2% of synapses are automatically linked to somata, limiting the relabeling to about 10% of the unlabeled cells, a substantially larger fraction of orphan axons—roughly 40%—were reassigned under this scheme (**Fig. 3C**).

To extend this analysis to additional unlabeled somata, we manually reconstructed axons for select cells in which no axonal reconstruction had previously been completed. **Fig. 3D** shows three such examples, each assigned to one of the three synapse polarity classes based on axon composition. When compared to their LM molecular labels, these polarity assignments corresponded to *vglut2a*-positive (green), *gad1b*-positive (magenta), and “other” (tan) cells, respectively. The tan neuron, identified as a serotonergic cell in the *raphe nucleus*, is consistent with a non-glutamatergic/GABAergic identity. Its axon exhibited a more even mix of excitatory-and inhibitory-appearing synapses. While this does not confirm that axons with mixed synapse predictions are modulatory, it justifies the inclusion of an “other” category by demonstrating that such synaptic patterns can correspond to known modulatory cell types.

The extensive set of re-labeled orphan axons and the broad coverage of input synapses on dendrites associated with somata allows us to assess the spatial distribution of synaptic input polarity across the brain, which we refer to as the “input drive index” (**Fig. 3E**). For this analysis we focused on the 41,000 neurons expressing *vglut2a* or *gad1b*, and distinguished between two input axes: excitatory versus inhibitory (E-I index, top row of mountain plots), and ‘fast-acting’ versus ‘other’ index (O index, bottom row). The results are robust to synapse threshold selection (**SFig. 6**), and reveal biological structure when compared to shuffled controls (**Fig. 3E**, **SFig. 7**).

Notably, although excitatory neurons numerically dominate the brain, the net balance of excitatory versus inhibitory axonal input is remarkably even. Theoretical and experimental studies have shown that maintaining a balance between excitation and inhibition is critical for stable, flexible neural computation, with disruptions linked to pathological activity or circuit instability ^37,38^.

While this balance is approximately maintained across the brain, there are striking exceptions. The optic tectum, for instance, exhibited a strong skew toward inhibitory input, while the habenula and cerebellum are skewed toward excitatory input drives. The excitatory bias in the habenula and cerebellum aligns well with prior anatomical and functional data. The habenula receives dense glutamatergic input from pallial sources, and the cerebellum is heavily innervated by glutamatergic mossy fibers and climbing fibers from the spinal cord, brainstem, and inferior olive.

In contrast, the inhibitory bias of the tectum was unexpected, as this region is typically thought to be driven by excitatory retinal ganglion cells (RGCs). We evaluated the relative contribution of RGC synapses to the 5 million synapses in the tectal neuropil and found that the total number of synapses in the retinal arborization fields, as estimated from the overlay to the zebrafish brain atlas, accounts for only 0.5 million. This indicates that RGC input constitutes only a small fraction of total tectal synapses, and suggests that the local tectal circuitry has an overall inhibitory bias, which is surprising.

We also asked whether excitatory and inhibitory neurons systematically differ in the net polarity of their synaptic input. At the coarsest level of anatomical resolution, we observed no significant difference between these two populations. However, when examining finer subdivisions (**Fig. 3E**), particularly within the hindbrain rhombomeres, clear differences emerged: inhibitory neurons tended to receive more excitatory input than their excitatory counterparts, and vice versa. This reciprocal arrangement is consistent with theoretical expectations for hindbrain central pattern generator circuits where cross-inhibitory and cross-excitatory motifs are used to generate rhythmic dynamics ^39^

Finally, across brain regions, we found that the relative contribution of the modulatory input, as defined by our O index, is consistent between excitatory and inhibitory neurons, suggesting that neuromodulatory inputs tend to co-regulate excitation and inhibition together, rather than selectively biasing one over the other. This further supports the idea that the E/I ratio is a conserved setpoint, maintained across brain states to ensure circuit stability.

### Inferring function from connectivity in the lateral line system

Larval zebrafish, like most teleosts, use the lateral line system to detect and interpret water flow across their body surface. This system is composed of mechanosensory organs called neuromasts, which extract directional flow information in four cardinal directions: headward (hwd), tailward (twd), upward (up), and downward (down) ^40,41^. Directional sensitivity is built into the system’s anatomy: the location of each neuromast on the animal’s body determines whether it responds to horizontal (hwd/twd) or vertical (up/down) flow (**Fig. 4A**, top: teal vs. vermillion), and within each neuromast, mirror-symmetric pairs of hair cells with opposing polarity allow discrimination between the two directions along a given axis—headward versus tailward or upward versus downward (**Fig. 4A**, bottom; golden brown vs. purple). Notably, headward and upward tuning, as well as tailward and downward tuning, have been shown to share common transcriptional control and can be grouped as polarity pairs ^42^. Together, a hair cell’s orientation and polarity define its *directional tuning*, making each cell a unique sensor for a specific cardinal flow direction.

We focused on the posterior lateral line (PLL), a major branch of the lateral line system that spans the trunk of the zebrafish. We used connectomics-based circuit analysis to inquire how the directionally tuned input of the lateral line is transformed by the hair cells’ first targets - the posterior lateral line ganglion (pLLg) and the downstream neurons in the medial octavolateralis nucleus (MON) (**Fig 4.C-D**). Specifically, we asked to what extent the four cardinal directions are maintained in “labeled lines”, and where and how they converge. Routing directional signals along labeled line pathways can bypass more elaborate computational processes, and enable rapid sensorimotor decisions essential for behaviors like escape or course correction ^43,44^. Convergence, on the other hand, could potentially enable the computation of higher-order flow features, such as net flow direction, flow divergence, or body tilt, which are relevant for behaviors such as rheotaxis, posture control, and escape ^45–48^.

We defined *orientation tuning* as sensitivity to either the horizontal (hwd/twd) or vertical (up/down) axis, and *polarity tuning* as the direction of flow within that axis - either positive (pol+, hwd or up) or negative (pol-, twd or down). Each hair cell’s individual preferred response direction is structurally encoded by the staircase-like arrangement of its stereocilia and the position of its kinocilium, such that flow in the direction of increasing cilia length drives depolarization ^49^. The preferred response direction of each hair cell can thus be inferred from its morphology and the four *directional tuning* vectors (hwd/twd/up/down) are defined by aligning these preferred vectors with the body’s major axes (black ellipses superimposed on hair cells in **Figure 4B**). It is helpful to compare this directional encoding to the tuning properties of direction selective ganglion cells in the vertebrate retina, which - in similar fashion - extract information about visual flow by responding best to motion in one of the same four cardinal directions (up = dorsal, down = ventral, hwd = nasal, twd = temporal)^50^.

As in the retina, the posterior lateral line system is organized symmetrically, with mirrored neuromast arrays and matching posterior lateral line ganglia (PLLg) on the left and right sides of the fish, providing an internal consistency control for connectivity analysis. Directional signals from hair cells are transmitted to afferent neurons in the PLLg, located near the ear on each side of the fish (**Fig. 4A–C**). These afferent neurons, referred to as PLLn, contact only hair cells of matching orientation and polarity—both within and across multiple neuromasts—resulting in the precise directional tuning of each individual neuron to one - and only one - of four flow directions (headward, tailward, upward, or downward; **Fig. 4C**) ^51,52^. In our dataset, each PLLg contains ∼75 mature, connected PLL neurons (79 on the left, 76 on the right), along with ∼50 immature afferents that have not yet formed significant postsynaptic connections and were excluded from downstream analyses (**SFig.8**).

Downstream of the PLLg, directional flow signals are processed by neurons in the medial octavolateralis nucleus (MON), a hindbrain region that integrates and relays this information for further computations and behavioral responses (**Fig. 4D**). To map how the unidirectional encoding of the PLLg is transformed into a combinatorial, analog representation of flow in the MON, we traced all ∼43,000 synapses from mature PLLg neurons to their postsynaptic targets. Approximately 40% of these synapses connected to somata of MON neurons, each of which received input from at least two different afferents—yielding ∼800 MONs on the left and ∼700 on the right. To characterize the directional tuning of each MON, we summed the contributions from its presynaptic PLL neurons and computed two indices: (1) an *orientation index* as the normalized difference between horizontal and vertical inputs, (2) *polarity index* as the normalized difference between a sensitivity toward positive (hwd/up) and negative (twd/down) flow.

Because each PLLn contacts only hair cells of a single type, both hair cells and PLLn neurons have a fixed directional tuning defined by a binary pair: orientation and polarity, with each value taking –1 or 1. In contrast, MONs are allowed to mix directional signals from the PLLns and as a result, their tuning can be described by two continuous values that together define an analog representation of flow direction.

While all MONs and their afferent input from the PLL are contained in the EM volume, the sensory axons of the PLLns and their associated hair cells were only partially captured. Specifically, only two neuromasts per side (D1 and OC1, **Fig. 4A-B**) fell within the reconstructed volume, allowing just 24 out of 155 PLL neurons (14 right, 10 left) to be directly traced to their presynaptic hair cells. These neuromasts included one with horizontal tuning (D1) and one with vertical tuning (OC1), enabling full assignment of orientation and polarity for this small subset. An additional ∼10 PLLn per side was partially labeled with orientation based on their projections to vertically tuned neuromasts (D2/D3; **Fig. 4A**), though their polarity remained unknown. The remaining ∼50 PLLn per side could not be labeled, as their presynaptic partners laid outside the volume (**Fig. 1C** - grey neurons).

We leveraged the partially labeled set of PLLn to infer orientation and polarity labels for the broader population of unlabeled PLLn and MON neurons using a bootstrap-based label propagation approach (see Methods). In each bootstrap iteration, balanced groups of labeled PLLns served as seeds, and labels were propagated through the synaptic network in repeated forward and backward steps. First, MON neurons received input from the seed groups, and a polarity or orientation index was computed based on the relative strength of those inputs. These MON indices were then backpropagated and used to update labels for connected PLLns. These steps were repeated in cycles, enabling label influence to flow bidirectionally through the network until assignments stabilized. To ensure robustness, 15% of synapses were randomly removed in each iteration. Final results were aggregated across 100 bootstrap runs to generate probabilistic maps of orientation and polarity tuning for the full MON and PLLn populations (**Fig**. **4E–F**; also see neuroglancer link <u>here</u>).

To evaluate our label propagation method, we compared the inferred PLLn labels against well-established anatomical and functional expectations. First, given the distribution of neuromast orientations along the trunk (**Fig. 4A** - top), we expected more horizontally than vertically tuned PLLn neurons. Second, because each neuromast contains roughly equal numbers of hair cells with opposing polarity, we anticipated a near-even split between pol+ and pol– afferents (**Fig. 4A** - bottom). Finally, due to the system’s bilateral symmetry, we expected similar results on both sides of the animal.

Our findings were consistent with these expectations (**Fig. 4E**, post-propagation rows). Initially, due to sampling bias, the orientation labels were skewed, with vertical flow directions overrepresented by ∼3:1 compared to horizontal (**Fig.4E**, pre-propagation rows, vermillion vs teal PLLn panel). After label propagation, this pattern reversed: we observed 39 horizontally tuned (teal) and 16 vertically tuned (vermillion) PLLn on the right, and 40 horizontal versus 6 vertical on the left. Polarity labels showed a near-even distribution throughout: 30 pol+ (golden brown) and 23 pol– (purple) on the right, and 36 pol+ and 31 pol– on the left. The small bias towards pol+ labels could either reflect an error in our propagation methods, or alternatively, it could be explained by a slight preference of pLLns for pol+ hair cells in the trunk neuromasts. All counts reflect afferent assignments which have been assigned to a category in more than 95% of the bootstrap iterations. These results, together with the overall conservation of mirror symmetry across both hemispheres, enhance our confidence that our label propagation method is both accurate and robust, capturing biologically plausible tuning distributions without overfitting to the sparse initial labels.

Finally, we asked how directionally tuned input from all afferent neurons is transformed by the MONs. Prior to label propagation, MON orientation and polarity indices appeared largely binary, reflecting non-overlapping input from a limited subset of labeled PLLn (**Fig. 4F**, pre-propagation histograms). Following propagation, these indices became more broadly distributed, consistent with convergence across PLLn inputs with diverse directional preferences (**Fig. 4F**, post-propagation histograms). This transformation preserved the original labeled-line axes at the corners of the tuning space while populating intermediate states, enabling both labeled line and integrated representations.

Notably, the post-propagation heatmaps exhibit a triangular density structure, indicating that not all combinations of preferred flow direction are equally represented. MON neurons receiving inputs exclusively from horizontally tuned neuromasts (right heatmap edge, orientation index=1) show a continuous representation of flow along the headward-tailward direction. On the other hand, MONs receiving inputs exclusively from vertically tuned neuromasts (left heatmap edge, orientation index =-1) preserve the labeled line structure of upward vs downward direction tuning. MONs near the center received mixed directional input, suggesting capacity for encoding net flow^46^ or responses to chemical cues like serotonin ^53^.

MONs receiving exclusively input from a given polarity tuning show a striking asymmetry: on the left side positively tuned inputs (upward and headward) are continuously represented, while on the right side the negatively tuned inputs (downward and tailward) are continuous. This results in an enrichment of headward-tuned MONs on the right and tailward-tuned MONs on the left side of the animal. Such cross-hemispheric lateralization may enhance sensitivity to counterclockwise flow around the body, a bias whose consistency across animals has yet to be investigated and whose functional significance remains unclear.

### The connectome of the hindbrain motion integrator

The zebrafish’s anterior hindbrain is home to a dedicated circuit structure that has been shown to integrate noisy visual motion and trigger targeted tail flicks once enough evidence is accumulated to pass a decision threshold ^54,55^. This circuit has two properties whose precise circuit implementations are not yet understood (**Fig. 5A**). First, it is assumed that the long integration time constants of several seconds are the result of a highly recurrent circuitry amongst dedicated excitatory and ipsilaterally projecting neurons. Second, swim direction is ultimately determined by an inter-hemispheric “winner-take-all” interaction, which is hypothesized to be implemented by long-range inhibition between the hemispheres. This inhibition could either be implemented through direct inhibitory contralateral projections, it could be communicated via excitatory interhemispheric crosstalk with specific inhibitory targets, or it could be mediated by an indirect pathway that is relayed through secondary nuclei. We set out to utilize our connectomic resource to test these hypotheses and thereby constrain and validate our existing circuit models ^54^.

To first obtain a small, unbiased sample of neurons in the integrator region of the right anterior hindbrain, we reconstructed a random set of 20 neurons in rhombomeres 1-3 (**Fig. 5A**, bottom), and found that 18 out of these cells could be divided, based on axonal projection patterns, into two separate classes: Class I neurons (7 out of 18) formed axons and dendrites locally in the ipsilateral hindbrain (**Fig. 5B**, top), which could constitute the anatomical substrate of a recurrent integrator circuit. By contrast, Class II neurons (11 out of 18) projected axons exclusively to the contralateral hemisphere and could thereby implement the hypothesized interhemispheric inhibition (**Fig. 5B**, bottom). In a companion paper ^56^, we developed a classifier trained on anatomical reconstructions of functionally identified cells and found that Class I and some Class II cells indeed corresponded with integrator neurons that were identified using functional imaging. In addition to these two main classes, we found two hindbrain neurons with stubby dendrites extending into the ipsilateral cerebellar neuropil whose axons projected to the ipsilateral cerebellar cortex (not shown). Because these cells had no synaptic connections to the rest of the circuit, and because we never identified this third cell class in the context of the coherent dot optomotor response stimulus in our companion paper, we excluded these two cells from further analysis.

We next set out to build a detailed and exhaustive reconstruction of the excitatory and inhibitory synaptic network that - within one hemisphere - gives rise to the integrator and “winner-take-all” circuit. To that end, we focused on the Class I cells, because - through their ipsilaterally projecting axons - only they are in a position to support the maintenance of long time constants through local recurrence. Class II cells cannot participate in this computation because they exclusively project contralaterally.

An initial analysis of the Class I axonal projection patterns revealed that 3 out of the 7 neurons projected and synapsed exclusively onto the dorsal raphe and did not obviously participate in local computations. We therefore limited our circuit tracing to the remaining 4 Class I seed cells and reconstructed their synaptic connectivity in depth across four network layers (**Fig. 5C**). Briefly, we reconstructed all presynaptic partners of these 4 seed cells (1st degree inputs, N = 68), all of their postsynaptic partners (1st degree outputs, N = 113) and all of the postsynaptic partners of all 1st degree outputs that were themselves Class I cells (2nd degree outputs, N = 825). The 2nd degree outputs were traced to determine if these cells had already been reconstructed in any of the other network layers - as a method to probe the level of recurrence in the anterior hindbrain. Indeed, we found that 89 of the 2nd degree outputs were already nodes in another network layer, whereas 724 cells were newly discovered. Interestingly, this means that 50% of neurons in the 1st degree input layer, seed cell layer, and 1st degree output layer were ‘rediscovered’ when tracing the 2nd degree output layer (i.e., 89 out of 179 individual neurons).

The resulting hindbrain network, reconstructed through both forward and backward tracing from four seed neurons, comprised 915 individual neurons and 1,652 manually annotated (ground-truth) synapses, corresponding to 1,332 unique connections, of which some were multisynaptic contacts. In summary, synaptic outputs were exhaustively reconstructed during the forward-tracing phase, in which all postsynaptic partners were identified for the four seed cells and first-degree output neurons classified as Class I. Backward tracing was similarly complete: all neurons presynaptic to the Class I seed cells were identified.

When we plotted all 1st degree inputs and 1st degree outputs (n = 181 cells) and color-coded these reconstructions by their neurotransmitter type (**Fig. 5D**), we found that glutamatergic neurons were clustered along the midline, but that their axons did not cross to the other hemisphere. By contrast, GABAergic neurons were located more laterally, and a large fraction of these projected contralaterally. Long-range inputs to the four Class I seed cells were ipsilaterally projecting, primarily glutamatergic and located in visual regions of the thalamus (n=8) and pretectum (n=5). Notably, we found that 60% of Class I cells were glutamatergic, and that Class II neurons were almost exclusively GABAergic (**Fig. 5E**).

This exhaustive multi-layer circuit reconstruction strategy (**Fig. 5C**) allowed for a detailed analysis of the recurrent hindbrain circuitry. To that end, we plotted all Class I and Class II cells within the 185 neurons that are composed of the seed cells, their inputs and their 1st degree outputs as a network graph (**Fig. 5F**). We found that all Class I cells were highly connected with each other via a ‘central integrator cluster’ that consisted of a strongly recurrent network of glutamatergic neurons with a smaller fraction of GABAergic neurons. Class II neurons, which are predominantly GABAergic, received strong excitatory input from the ‘central integrator cluster’ and projected exclusively to the other hemisphere.

For a quantitative analysis of these hindbrain connectivity motifs, we plotted the exhaustively reconstructed input connectivity diagram of the four Class I seed cells (**Fig. 5G**) and the output connectivity diagram (**Fig. 5H**) of all Class I cells for which we had exhaustively reconstructed outputs (seed cells and 1st degree outputs). While Class I cells received a strong excitatory drive from long-range inputs originating in the pretectum and tectum, their strongest excitatory inputs in fact originated from other Class I neurons in the hindbrain (**Fig. 5G**). In accordance with this, we found that Class I neurons formed a large fraction of their output synapses with other Class I cells, and - to an even stronger degree - with Class II neurons in the same hemisphere (**Fig. 5H**). Interestingly, we also found excitatory outputs onto a population of neurons whose axons project posterior, where they could innervate spinal projection neurons, which could trigger a motor command that leads to targeted tail flicks once enough evidence is accumulated.

In summary, our analyses confirmed our hypothesis that the hindbrain contains a highly recurrent excitatory network of Class I neurons that receive long-range excitatory inputs from ipsilateral visual regions. This finding makes the alternative option, where recurrency is implemented through relays via alternate brain areas, less likely. The integrated information is sent - via long-range inhibition mediated by a population of GABAergic Class II neurons-to the contralateral hindbrain, where it provides graded inhibition that can implement an inter-hemispheric “winner-take-all” mechanism by which the certainty level of a binary decision can be encoded (**Fig. 5I**).

### Characterization of the synaptic outputs of a tegmental excitatory nucleus (TEN) reveals a potential source for brain-wide neuromodulation

We next wanted to leverage connectomics to contextualize the function of a group of glutamatergic cells in the tegmentum that we refer to here as the Tegmental Excitatory Nucleus (TEN). The TEN, which contains sparse expression of the neuropeptide cocaine-and amphetamine-regulated transcript 2 (*cart2*) ^28,57^, is likely homologous to the mammalian Edinger Westphal nucleus, a subregion of the mammalian periaqueductal gray (PAG), which is involved in regulating the behavioral and autonomic responses to arousing and threatening stimuli ^58,59^. In fish, functional studies of neurons in this area have identified correlations with arousal motor behavioral states and cardiac activity during threat-related challenges ^57,60^. These correlations single out this region as a potential node that integrates external information, singles out appropriate efferent targets - skeletal or visceral - and modulates internal states accordingly. To anatomically constrain the role of the TEN in regulating either motor actions, visceral actions, or both, we harnessed our connectomics resource to identify its specific target neurons.

Accordingly, we first identified this region in the EM volume based on glutamatergic fiduciaries, which form an isolated island around the ventricle that is just ventral to the optic tectum, and reconstructed a subset (*n = 111* out of 191 cells) of the neurons whose somata reside in this volume (**Fig. 6A**). We found that the dendrites of these cells were confined to nearby locations in the thalamus, while the axons distributed widely - collectively spanning nearly the entire hindbrain (**Fig. 6B**). Based on these axonal projection patterns, we defined four morphological classes (**Fig. 6B,C**): 1) axons that remain within the midbrain, 2) axons that project ipsilaterally along the midline into the inferior olive, 3) other ipsilateral projections that distribute more widely across the hindbrain, and 4) axons that cross the midline in the thalamus and then turn posteriorly into the hindbrain where they ultimately reach the dorsal-posterior brainstem in the vagal neuropil region.

To identify post-synaptic partner cells, we used the automatic synapse detection and applied a synapse volume threshold (0.03 *μm*^3^) to reduce the false-positive rate, yielding 5297 synapses. From these synapses, 40.1% of the postsynaptic dendrites were associated with a soma, yielding 778 unique neurons (**Fig. 6D**). Consistent with the diversity of projection morphologies, we found that these four classes innervate distinct populations of neurons, where only 7 out of 778 targets were innervated by multiple classes. Class 1 TEN neurons exclusively connected with other neurons in the midbrain, particularly cells lateral to the TEN near the nMLF and oculomotor nuclei. On the other hand, we saw that both Class 2 and 3 neurons target cell types implicated in regulating different aspects of behavioral control, such as motor adaptation via climbing fibers of the inferior olive (innervated by Class 2) ^61^, Mauthner cell excitability via spiral fiber neurons (innervated by Class 3) ^43^ and coordinating spinal cord central pattern generators via reticulospinal projection neurons (also innervated by Class 3). The outputs of Class 4 neurons also contained premotor neurons, including reticulospinal projection neurons and spiral fiber neurons. There were also many postsynaptic neurons in the dorsal brainstem with functional roles that are less clear. However, we observed that the axons of Class 4 neurons pass through the neuropil associated with the vagal system and weave remarkably close to the fascicules of outgoing vagal motor neurons (**Figure 6E**).

The observed proximity of TEN axons and vagal motor nerves suggests that Class 4 TEN neurons modulate vagal motor nerve activity - a hypothesis that could explain the previously described temporal correlations of TEN activity with heart rate. However, we found that all of the identified targets of the TEN neurons were interneurons with axons contained within the brain, and thus were not part of the organ-innervating motor vagus system. To test whether there is an indirect connection between the TEN neurons and the vagus, we first reconstructed a large number of motor vagal neurons (*n = 393,* over 60% of 631 identified motor axons) - which we identified by tagging axons that travel out of the body through the right vagus nerve and tracing back dorsally to their cell body in the brain ^62^. We then reconstructed a sparse set of presynaptic inputs to these cells (*n = 139*) to look for any connectivity with the outputs of type IV TEN cells. While the somata of cells downstream of the TEN and those related to the vagus motor control were intermingled and occasionally direct neighbors, we identified no true synapses (5 detected “synapses” were false positives) between these populations (**Fig. 6F**). This connectomic separation emerges from the anatomy of the neuropil, where we observed 3 distinct zones, with two vagal related layers surrounding one occupied by TEN related axons (see neuroglancer link <u>here</u>). Collectively, these data support a role for the TEN in modulating motor output, while allowing us to reject the hypothesis that they - directly or indirectly - govern vagal motor activity through synaptic transmission (**Fig. 6G**).

## Discussion

The dataset presented here provides a structurally rich, multimodal view of the larval zebrafish brain. Although the EM volume is not a complete connectome, with only approximately 20% of total neurite path length assigned to somata, it nonetheless enables foundational analyses. By integrating ultrastructural data with light microscopy-based labeling of glutamatergic and GABAergic neurons in the same animal, we constructed a comprehensive map of excitatory and inhibitory output balance across all brain regions. This analysis revealed that these molecularly defined classes represent only a fraction of the total cellular population in each area, a distinction that could not be resolved by light microscopy alone, where unlabeled cells remain invisible.

The structural regularity of the zebrafish brain enabled generalizations from limited ground truth, allowing us to scale molecular annotations across millions of synapses. Specifically, we trained a synapse classifier on a subset of axons with known cell bodies, which we then applied across all ∼30 million axonal-dendritic synapses. Using a Bayesian inference framework that applies Dale’s law to axonal fragments with multiple outputs, we assigned polarity to approximately 40 percent of all axon fragments. We found that more than 80 percent of the ∼110,000 soma-associated dendrites received input from these classified axons, enabling us to compute regional input indices that quantify how excitatory, inhibitory, and modulatory signals shape local circuit dynamics.

Similarly, we used the regularity of synaptic connectivity to infer the functional tuning of lateral line sensory neurons, even with incomplete peripheral tracing. Our analysis shows that detailed anatomical data can reveal new principles of vertebrate circuit organization. In the lateral line, we find that directional signals, initially segregated along labeled lines, are transformed into distributed representations in the hindbrain. The surprising hemisphere-specific asymmetries in tuning can either be due to stochastic variation during development or could be a stable zebrafish phenotype like the strong asymmetry observed in the habenula. The hypothesis of stochastic developmental variability can be explicitly tested by repeating the analysis in other animals, where the asymmetry might be inverted, absent or remain the same. Future comparative and functional studies will be essential to uncover the origins and adaptive significance of these asymmetries.

Rigorous experimental design demands that hypotheses be framed in ways that allow for explicit falsification ^63^. While supportive evidence can increase confidence, it is the ability to disprove a model that carries the greatest weight ^64^. Connectomics-driven tests of circuit models preserve this principle: by predicting specific anatomical motifs they provide clear criteria for falsification - if the motifs are absent, the model is invalidated.

Notably, we highlight the negating power of connectomics by rejecting a hypothesis that TEN neurons modulate heart rate via direct input to the motor vagus ^57,60^. Despite their anatomical proximity, we found no synaptic connections—direct or indirect—between TEN neurons and vagal motor populations. Instead, TEN outputs target premotor circuits involved in locomotion, including reticulospinal and inferior olive-projecting neurons. These findings highlight how high-resolution connectivity can falsify plausible models, refining our understanding of brain-wide control of internal state.

On the other hand, the hindbrain motion integrator provides a complementary example in which reconstruction supports a mechanistic model - but crucially, one that could have been falsified. Based on prior behavioral and physiological evidence, it was hypothesized that sustained activity in this circuit arises from local excitatory recurrence and is shaped by inter-hemispheric inhibition^54^. We identified a densely interconnected group of glutamatergic neurons that form a recurrent excitatory network and receive excitatory input from visual regions in the thalamus and pretectum, suggesting a role in accumulating sensory evidence. In turn, they drive GABAergic neurons that project to the opposite hemisphere, forming a “winner-take-all” circuit that converts graded input into binary motor decisions. This connectivity supports a decision-making mechanism in which local excitation integrates sensory input over time, and long-range inhibition encodes the chosen outcome.

Beyond the examples discussed here, our multimodal dataset expands the space of exploratory and hypothesis-driven studies of vertebrate brain circuitry. The segmented fragments are large, proofreading is tractable, and all synapses are annotated, which offers a realistic pathway to targeted reconstruction. Indeed, our semi-automated workflows dramatically reduce the labor required to reconstruct single neurons, with typical efforts requiring just 10 minutes to 1 hour per neuron, compared to days in unsegmented datasets. This shift lowers the barrier to circuit dissection, enabling broader community access to hypothesis-driven connectomics. Still, limitations remain. Many modulatory cell types are not yet fully reconstructed, and significant axonal tracing is required to extract full connectivity matrices. Addressing these challenges through community curation, collaborative proofreading, and continued annotation will further enhance the coverage and reach of this resource.

## Acknowledgments

1. F. Engert received funding from the National Institutes of Health (U19NS104653 and 1R01NS124017-01), DoD (W911NF2420112) and the Simons Foundation (SCGB 542973 and NC-GB-CULM-00003241-02. J. W. Lichtman received funding from the National Institutes of Health (U24NS109102, U19NS104653, and UM1NS132250). This project was partially supported by NIH grants 1U01NS132158 and R01HD104969 to H. Pfister, Simons Foundation SCGB 542943SPI to M. Ahrens and by the Howard Hughes Medical Institute.

We are grateful to F. Collman, D. Brittain, A. Halageri, and S. Dorkenwald for constructive feedback and advice on the CAVE deployment. P. Petkova contributed critical segmentation of both LM and EM datasets and generated manual segmentation ground truth, for which we are especially grateful. We thank V. Wang and A. Chen for providing example neurons used to test the polarity classifier. We also acknowledge B.C. Oban and T. Skye for maintaining a supportive working environment. Finally, we thank J. Miller and K. Hurley for their expert care and management of the fish facility, and E. Nako and S. Montgomery for their administrative support.

## Author contributions

Conceptualization: M.D.P., M.J., K.J.H., G.F.P.S., F.E., J.W.L.; LM data acquisition: M.D.P.; Fish line generation: J.B.W.; EM data acquisition: M.D.P., R.S., J.B.W., J.C.T, N.I.; Image stitching, alignment, normalization: A.P., S.W., Y.W., M.J., T.B., M.D.P.; CLEM: M.D.P., Z.T.M.; Atlas registration: M.D.P., S.K.V., A.B.; Ground truth generation for segmentation models M.D.P., K.B.H., M.J., E.C.P, M.T., Z.D., S.D., Z.F., K.I.K., P.G., J.W-C., D.A., M.S.V., T.T.C., S.M., T.Y., A.D., J.M.D-M., S.B., N.S-M.,E.G., M.C., S. F-M., M.A.F., G.P.H., J.K., M.L., A.L., C.O., A.L.S., S.T., C.W., J.J.W.; Ground truth generation for synapse models: M.D.P., K.B.H., S.C., A.V.M., D.C., E.C.P., X. L., J.W-C; Synapse model training: T.B., D.W., Z.L.; Segmentation model training: M.J.; Model performance validation: M.D.P., M.J.; Automated segmentation and synapse analysis: M.D.P, M.J.; Input/output polarity drive index analysis: M.D.P., M.J, F.E.; Lateral line afferent tracing and analysis: M.D.P., M.J., F.E., J.W-C.; TEN neuron tracing and analysis: K.J.H; Hindbrain integrator neuron tracing and analysis: G.F.P.S., R.T., M.S., D.H., A.H., S.D., R.C.W., L.L.Z.; CAVE deployment and maintenance: J.C.,J.T.; VAST integration: D.R.B, E.C.P.; Project supervision: M.D.P., V.J., F.E., J.W.L, S.B., C.K, W.K., G.W.M., M.B.A, H.P.; Manuscript writing: M.D.P., M.J., K.J.H., G.F.P.S., F.E.; Correspondence: M.D.P., F.E., M.J., J.W.L

## Data availability

All proofreading and segmentation data are accessible through the CAVE platform, which supports collaborative editing, programmatic access to annotations and meshes, and a changelog of proofreading history. Individual galleries on the project <u>website</u> showcase the specific research vignettes explored in this paper, allowing direct visual inspection of the underlying data.

## Supplementary material

### Animal care

To label the major neurotransmitter types and the vasculature we crossed *Tg(gad1b:GFP)* with *Tg(flk1:mCherryCAAX;vglut2a:loxP-DsRed-loxP-GFP)* \ in Nacre (transparent) background ^65^. Individual *Tg(vglut2a:loxP-DsRed-loxP-GFP)* and *Tg(gad1b:GFP)* lines were previously generated by BAC transgenesis ^66^. These lines label the majority of excitatory and inhibitory neuronal subtypes in the optic tectum and hindbrain circuitry. The *Tg(flk1:mCherryCAAX)* was previously generated using classical transgenesis by cloning of the endothelial-specific *flk1* promoter ^67^.

Brain tissue from rats which had been sacrificed for other experiments served as support tissue for zebrafish embedding.

### Confocal image acquisition

At 7-day post fertilization a larval zebrafish was immobilized in 1.8% low-melting-temperature agarose in a glass bottom dish and imaged with a ZEISS LSM 880 confocal microscope with W Plan-Apochromat 20x/1.0 Korr DIC M27 75mm objective. The dataset encompasses the entire brain (2 overlapping tiles, 607×607×317 μm, 0.506×0.506×0.652 μm³ voxel size) and consists of three 16-bit channels: (1) vasculature and excitatory neurons *Tg(flk1:mCherryCAAX;vglut2a:loxP-DsRed-loxP-GFP)* excited with a DPSS 561-nm laser, (2) inhibitory neurons *Tg(gad1b:GFP)* excited with an 488-nm argon line, (3) transmitted light.

### Sample preparation for electron microscopy

Following confocal imaging, the agarose-embedded zebrafish was anesthetized with 0.02% (w/v) tricaine mesylate (MS-222, Sigma-Aldrich) and dissected as described previously ^68^. The brain was exposed and the tail was removed just anterior to the caudal fin for notochord access. The fish was fixed using microwave irradiation series ^69^ in 2% glutaraldehyde, 6% mannitol solution, followed by overnight incubation in fresh fixative at 4⁰C. A modified rOTO protocol ^70^ was used for parallel staining of the fish and rat support tissue for electron microscopy. After sodium hydrosulfite reduction ^71^ and osmium incubation, the samples were reduced with potassium ferrocyanide and stained with thiocarbohydrazide, osmium and uranyl acetate. Dehydration with ethanol and infiltration with low-viscosity LX-112 resin preceded embedding with the fish oriented within a rat tissue support block (2mm x 2mm x 3mm with 0.75mm punch hole to house the fish). The block was cured at 60⁰C for 75hrs. The detailed protocol with reagent information, solutions and incubation times is provided as <u>Supplementary File #1</u>.

### Sample sectioning and wafer fabrication for electron microscopy

The resin block was trimmed to a coffin shape measuring 4.2mm length x 1.6mm width using a Diatome Trim 90 diamond knife (Diatome, USA) and ultramicrotome (UC6, Leica, Germany). The leading and trailing edges were trimmed at 100⁰ and 60⁰ angles, respectively. The fish was cut from ventral to dorsal direction in 30 nm thick sections using a series of five Diatome Ultra 45/Ultra 35 diamond knives. This process was performed over 4.5 days using an automated tape collection ultramicrotome (ATUM) system ^72^ set at a cutting speed of 0.3mm/s, resulting in a collection of 17,500 sections on 110 meters of carbon-coated Kapton tape. The tape was cut into strips and mounted on 126 square silicon wafers (University Wafers, USA) using double-sided carbon tape (Ted Pella, USA). The wafers were post-stained with uranyl acetate and lead citrate, and stored for imaging with a multibeam scanning electron microscope (mSEM, Zeiss) following the procedure in ^2^.

### Electron microscopy image acquisition

In preparation for imaging with the multibeam scanning electron microscope (mSEM), a Zeiss reflected light microscope (Axio Imager) was utilized to generate a high-resolution optical image (2.5 μm/px) of the entire wafer. This image served as a map to guide the automated acquisition process with a Zeiss mSEM equipped with 61 electron beams. The fish volume was imaged in two phases (1) 6,332 sections containing approximately 80% of the brain, and the total amount of data acquired was 350 TB, (2) 1931 sections, comprising the remaining dorsal portion of the brain, approximately 10% of the brain, amounting to 20 TB of image data. A total of 213 sections in phase1, and 174 sections in phase2 were skipped during cutting, leading to double-thickness sections (60 nm). It is important to emphasize that the cutting process never resulted in consecutive stretches of three or more damaged or missing sections. Additionally, a total of 39 sections across both phases were excluded from imaging due to severe cutting damage. The image acquisition parameters were set at 4 nm/pixel with a landing energy of 1.5 kV. Dwell times of 200 ns and 400 ns were employed for phases 1 and 2, respectively. The increased dwell time in phase 2 aimed to enhance the signal-to-noise ratio (SNR) due to the lower contrast arising from reagent washout at the sample interface.

### msem manager

We developed a custom workflow management software for Zeiss MultiSEM to run in the background during image acquisition. The software constantly monitored the data outflow from the microscope, and immediately sent email alerts to microscope operators upon detecting unexpected workflow interruptions. Once the microscope finished acquiring data for each section, the software also immediately started a series of data validation steps, including assays of data completeness, a stitchability test to ensure sufficient overlap between neighboring image tiles, detection for stage-settling vibration, identification of significant image distortion, and evaluation of region-of-interest targeting. In addition to these background data validation routines, this software also provided a visualization tool enabling the operator to intuitively identify out-of-focus images. Data validation results, along with the user input from the focus examination, were automatically compiled into an interactive HTML report, assisting the operator to promptly make retake decisions. The software is available on GitHub (https://github.com/YuelongWu/mSEM_workflow_manager).

### Stitching and full resolution tile rendering

Rigid stitching was performed on the raw data coming from the mbeam microscope for phase 1 and most of the sections of phase 2. Firstly, all the overlapped raw tile pairs were found based on the coordinates of each tile output by the microscope. Then SIFT features were extracted from the boundary area of each pair of overlapped raw tiles and all the tiles in each section were thus matched. For the tiles with too few SIFT features that failed to be matched (such as the resin regions), we estimated the rigid transforms for those tiles based on the transform of neighborhood tiles. Finally, a global optimization step made the matching results smooth through the whole section. Stitching of phase 1 was done on two Intel servers, each of which has 224 CPU cores. Stitching of phase 2 and full resolution tile rendering of both phase 1 and phase 2 was done on a few Google virtual machines, each of which has 96 CPUs. The stitched sections were rendered at full resolution and each section was cut into 4096*4096.png tiles without overlap between tiles. The rendered, stitched sections were the basis of the following alignment step. No contrast enhancement techniques such as contrast-limited adaptive histogram equalization (CLAHE) was used in this rendering step.

### Thumbnail alignment

We introduced low-resolution thumbnails to register the sections roughly in place in preparation for a fine alignment step. To generate the thumbnails, the full-resolution sections were first downsampled to a 64 nm pixel size. This was followed by bandpass filtering to equalize brightness variations and enhance membrane features in the raw data. Subsequently, the images were downsampled to their final 512 nm thumbnail pixel size. The thumbnails generation happened in real-time with image acquisition on an 8-core Windows desktop connected to the microscope buffer storage. The thumbnail of each section was matched with its ±2 neighbors. Initially, a feature-based method was used to estimate an affine transformation. This was followed by template matching on small, evenly distributed image patches on the sections to obtain corresponding points for more refined nonlinear deformations. Other than immediate neighbors, we also selected some “key frame” sections, on average 17 sections apart from each other, and matched them among themselves. These long-distance matches helped to minimize the z-direction drift caused by the accumulation of small alignment inaccuracies. We then modeled each section as a spring mesh (equilateral triangle mesh with a 25 μm grid size) and used the corresponding points found in the matching step to drive the meshes in a mesh-relaxation process and thereby obtain the alignment transformations. The matching and the mesh-relaxation of the thumbnail alignment step took about a day on an 8-core Windows desktop. These transformations were upsampled and applied to the rendered, stitched images at full resolution, which served as the input for the SOFIMA alignment step.

### High resolution EM alignment

We used SOFIMA ^73^ to obtain a precise alignment of the complete stack, following the procedures outlined previously ^2^, with some dataset-specific adjustments.

We used z = 2150 as the “anchor” section, and optimized the stack independently towards lower and higher z coordinates. Similarly to prior work, we used patches of size 160×160 pixels and a stride of 40×40 pixels for flow estimation. Flow fields were computed using 8×8, 16×16, and 32×32 nm^2 pixel images only given the high quality of the initial low-resolution alignment. We used a manually painted low-resolution “brain mask” to avoid computing flows and optimizing meshes in areas containing no tissue of interest.

Visual inspection of the aligned volume revealed remaining distortions around folds in the form of tissue “pinching”, which we seeked to minimize by running SOFIMA again using a higher resolution mesh and flow field (with a stride of 10×10 pixels, necessary to resolve the higher flow gradients around wrinkles) and restricting calculations to regions that our heuristics detected as likely to benefit from further corrections (where the residual flow magnitude exceeded 20 nm).

### Correlative light and electron microscopy

To register confocal microscopy (LM) and electron microscopy (EM) volumes of the zebrafish brain both volumes were resampled to isotropic resolution (EM: 0.512 µm x 0.512 µm x ∼0.480 µm; LM: 0.65 x 0.65 x 0.65 µm³), and segmented to produce a list of cell body locations (EM: 187,053 cells; LM: 10,937 *gad1b* cells and 24,812 *vglut2a* cells, **SFig.1A**). Excitatory and inhibitory neurons were segmented from confocal image stacks using a machine learning-based workflow (Ilastik) to classify 3D voxels for each channel, generating binary masks that were manually refined in VAST. Individual cells were then segmented using distance and watershed transforms in MATLAB, followed by manual verification of centroids to confirm correspondence with cell bodies, discarding ambiguous cases due to dense packing. For electron microscopy images, a custom MATLAB script was used to enhance cell-like structures through directional erosion, intensity adjustment, and morphological filtering. The resulting binary mask was manually corrected in VAST and segmented using distance and watershed transforms in MATLAB. This volumetric instance segmentation was inspected and further refined in VAST.

We used BigWarp ^74^ for initial landmark-based alignment with ∼1000 manually identified points, followed by iterative point cloud-based refinement of the alignment (**SFig.1B**). A custom MATLAB algorithm using Iterative Closest Point (ICP) identified corresponding cell groups between LM and EM data, generating a list of putative matches filtered based on distance, rotation, translation, and occurrence frequency. Specifically, groups of 3-8 closest LM cells were considered, and ICP found matching configurations among the 20 neighboring EM cells. The resulting putative matches were filtered: only matches with an average root mean square error of 0.5-1 µm, rotation < 0.1 radians, translation < 5-10 µm, and matches which occur at least twice were considered good. These thresholds were empirically determined to avoid ambiguities. Matches became landmarks for generating a new transform via a custom MATLAB driver for BigWarp, and the process iterated until no new matches were found. Manual intervention with additional landmarks was possible in poorly matched regions. This iterative procedure yielded 9,421 landmark points achieving single-cell matching across most of the brain (**SFig.1C-D**). All manual landmark points are dropped in the process, and the final landmark point list consists of landmarks that have passed the threshold selection criteria. The LM volume was registered to the Z-brain reference atlas using ANTs ^75^, and we combined the transforms to map the Z-Brain reference atlas to the EM volume for additional context.

To further evaluate the registration, the LM volume was mapped to the EM volume with a thin-plate spline transform calculated from the 9,421 landmarks and the registration at each of the 35,749 LM cell centers was evaluated by a human expert. For the evaluation, the human was presented with 20 µm x 20 µm image overlays between the EM and each of the two channels (*gad1b* and *vglut2a)* centered at the LM cell location: the registration was considered a match if every cell in the field of view from both channels was unambiguously assigned to an EM counterpart. We found that registration quality progressively decreases with distance from landmarks (**SFig. 2A**) and determined that cells located as third neighbors or further (10 μm or more from the landmark center) are predominantly non-matches. Most of the LM cells fall within the 10 µm radius from a landmark and are well-matched; there are 24,462 cells within 10 µm radius, 21,303 cells labeled as matches by secondary inspection. The remaining 10,045 cells further than 10 µm from a landmark are predominantly located in the edges of the volume and 1,479 cells are labeled as matches by secondary inspection (**SFig. 2B**).

The instance segmentation of cells in the LM confocal images was conservative, leading to undercounting of the labeled cells and assigning fewer labels to the EM cells after the registration of the two volumes. To rectify this issue, we implemented an intensity-based label assignment to the EM cells in the registered regions. There are 90,143 EM cells within 10 µm of a landmark, which we sorted based on the average intensity from the individual LM channels within the cell volume. We evaluated cells based on 20 µm x 20 µm image overlays and assigned them as *vglut2a* or *gad1b* based on signal from the LM images, or called them unassigned if there was no signal within the EM cell (**SFig3.A**). We evaluated 65,296 EM cells, assigning 26,117 *vglut2a,* 13,966 *gad1b* and 25,213 unlabeled cells. We observed labeled cells in the remaining 24,847 EM cells from the list at a very low rate, so we defaulted these to unlabeled without explicit screening. Additionally, we evaluated 5, 903 EM cells further than 10 µm of a landmark, assigning 798 *vglut2a,* 544 *gad1b* and 4,561 unlabeled cells. The distribution of labeled cell density in the volume is shown in **SFig.3B**, and the counts of *vglut2a* and *gad1b* cells relative to all cells is shown in **SFig.4A-B** for the major brain regions. Most of the assigned cells are immediate neighbors to the landmarks (**SFig.4C**). Finally, we defined a label index (I-E)/(I+E), where I and E are the fraction of cells which are *gad1b* and *vglut2a* within a column of 5 µm x 5 µm extension in xy, and 45 µm in z. This index quantifies the relative ratio of *gad1b* and *vglut2a* and mapped the spatial distribution of relative abundance for the two labels (**SFig.4D**).

Finally, we can use this data to estimate the total number of *vglut2a*-and *gad1b*-expressing cells in the brain. A total of 24,749 neurons from the LM point-cloud segmentation fell within 10 µm of an EM–LM landmark, the spatial range over which we retained high confidence in cross-modal matches. Within this same region, our exhaustive fluorescence-overlay evaluation identified 26,117 *vglut2a*-positive and 13,966 *gad1b*-positive neurons, for a total of 40,083 labeled cells. This indicates that our conservative LM segmentation approach recovered approximately 62% of the neurotransmitter-labeled population in the well-aligned volume.

The remaining 11,000 LM points lie outside the high-confidence matching radius. Assuming that LM undersegmentation is spatially uniform and continues to recover approximately 62% of labeled cells in these outer regions, we estimate that an additional ∼17,740 labeled neurons were missed. This brings the total number of *vglut2a*-and *gad1b*-expressing cells in the brain to approximately 58,000.

### Instance segmentation models

We collected instance segmentation ground truth within densely annotated subvolumes, either through de-novo manual voxel painting in VAST ^32^ or voxel-level corrections to automated segmentations generated with a machine learning model. In total, 10,990 μm^3^ were annotated in soma-dense regions (across seven 16.4×16.4×6.0 μm^3^ subvolumes, 40-80h manual effort per subvolume), and 1,378 μm^3^ in neuropil-dense regions (across seven subvolumes; six 4×4×8 μm^3^ subvolumes with 100-150 hours of manual voxel painting per subvolume, and one 6.1×10×10 μm^3^ subvolume with manual corrections of automatically generated segmentation).

We used this data to train three separate Flood-Filling Network (FFN) models, operating on 32×32×30 nm^3^, 16×16×30 nm^3^ and 8×8×30 nm^3^ images (henceforth called “32 nm”, “16 nm”, and “8 nm” models, respectively). All models used the annotations in the soma-dense regions, but only the 16 nm and 8 nm models used the annotations in the neuropil-dense regions. The FFN models used an architecture and settings identical to prior work ^29^, but the 32 nm model had an enhanced field of view of 33×33×33 voxels (vx), uniform 8 pixel step size in the in-plane and axial directions, and 16 residual modules. Unlike prior work, the centers of the training examples for the FFN were formed by taking the nodes of skeletons automatically ^76^ created from the volumetric annotations, post processed to maintain an average node separation of 300 nm and distance from the endpoints of at least 100 nm.

### Ouroboros

We applied the FFN segmentation models as described above to four 16×16×15.9 µm^3 subvolumes and four 16×16×10.2 µm^3 subvolumes of another larval zebrafish dataset acquired with a setup similar to the dataset acquired here. The subvolumes were targeted towards dense neuropil regions and their locations were selected to maximize diversity of the covered tissues. We then exhaustively proofread all neurites within these subvolumes and trained updated FFN models with this data. We kept the overall architecture and training procedures unchanged, but extended the network depth to 20 residual modules and used the AdamW optimizer, and trained a single variant of the network, operating on 16×16×30 nm^3-sized voxels.

We found these updated models to generate higher quality segmentations within the dataset from which the training data was extracted. We hypothesized that the gains in segmentation accuracy might transfer back to this dataset (“ouroboros”). We confirmed this hypothesis by segmenting the whole volume de-novo, starting with the existing, low-resolution (32×32×30 nm^3) soma-only reconstruction at the initial state of the new segmentation.

While the 2.5x increased network capacity and an order of magnitude increase in the amount of training data resulted in significant gains in reconstruction accuracy, we observed an increased rate of false merge errors in proximity of tissue folds, which were largely absent in the ground truth subvolumes the network was trained on. To mitigate this problem, we computed the oversegmentation consensus between the original and ouroboros segmentations. We then built a new agglomeration graph combining the agglomeration edges generated with the original and ouroboros models, and added edges reassembling the consensus supervoxels back to their shapes from the ouroboros segmentation. This moved any new merge errors from the supervoxel level to the agglomeration level, where they remain fixable through manual edits in CAVE.

### Semantic segmentation models

We trained convolutional neural networks to perform seven separate semantic segmentation (voxel-wise classification) tasks: two multiclass (tissue type classification, subcompartment classification) and five binary (myelin, glia, extracellular space (ECS), non-neural tissue and artifact detection). Training examples were always sampled at equal frequencies for all classes. They were extracted from the skeleton nodes of automated skeletons generated from test instance segmentations of the volume or from densely annotated instance ground truth subvolumes, except for the tissue type classification model and artifact detection models, which used manually painted voxel annotations.

The default architecture of these neural networks was a residual 3d convolution stack (same as the FFN), with a depth of 8 residual modules, a field of view of 33×33×33 vx, 3×3×3 convolutions using’valid’ mode and 32 feature maps, and residual summation discarding context at the edges of the larger-sized argument. Two exceptions from this default setup were made: the subcompartment classification model had a depth of 16 residual modules and a field of view of 65×65×65 vx, and the artifact detection model used a convolution-pooling network as in prior work ^29^.

We used three variants of the non-tissue detection model, with 32×32×30 nm^3^, 64×64×60 nm^3^, 128×128×120 nm^3^ input images and points extracted from 18,796 segments (3.4M tissue, 75.5M non-tissue). The artifact detection model used 16×16×30 nm input data and (artifact: 65.4 Mvx, regular data: 236.6 Mvx). The tissue classification model used 32×32×30 nm input data and annotations from the following classes: neuropil (61.5 Mvx), do not segment (167.2 Mvx), cell bodies (121.1 Mvx), folds (572 kvx), hair cells (1.3 Mvx), non-neuron cells (9.5 Mvx), support tissue (51.1 Mvx).

The remaining models used 16×16×30 nm input data. The subcompartment classification model used points from 165,015 segments across 3 classes: axon (2.4M), dendrite (94k), glia (985k). The binary classification models used the following annotations: extracellular space detection (8,287 segments; ecs: 56k, not-ecs: 301k), myelin (8,199 segments; myelin: 111k, not-myelin: 301k), glia (1,451 segments; glia: 5.7M, not-glia: 172k).

Instance segmentation assembly

We used the FFN models to first build a “base segmentation” optimized to reduce the frequency of merge errors. This was followed by an agglomeration procedure which reduced split errors while keeping merge errors manageable.

The section range 5333 <= z <= 5348 was a “coming-in” region bridging two parts of the volume cut with different diamond knives and with the original blocks oriented at a slight angle, such that the sections became progressively larger as they were sequentially cut. At all stages of segmentation, if a section had no tissue content (as determined by the section mask) locally within the current 500×500×500 vx subvolume, it was completely omitted from processing, and the corresponding segmentation was generated by duplicating content from the preceding section.

### Neuron fragment agglomeration

We excluded from agglomeration any segment that did not have a dominant classification (by fraction of labeled voxels) as “soma” by the tissue type model, and was classified as at least one of: 1) “not tissue” by the 32 nm or 64 nm non-tissue detection model, 2) “do not segment” or “hair cells” by the tissue type model, 3) “glia” by the glia detection model, 4) “extracellular space” by the ECS detection model, 5) “myelin” by the myelin detection model. Within phase2, we did not use the myelin classification as we found that the underlying model did not generalize sufficiently well to that part of that volume. Similarly, a minimum segment size of 100k voxels was additionally used for the conditions 3 and 4 above in phase2.

During agglomeration graph assembly, edges were processed in order of an agglomeration score as explained above, and soma separation was maintained by discarding any edge that would cause two known soma segments to be part of the same connected component. Similarly, we kept track of the fraction of voxels of every agglomerated segment classified by the subcompartment model as 1) axon or dendrite, 2) glia. When the classified voxel count exceeded 10,000 and the fraction associated with one of two options exceeded 0.75, the whole connected component was labeled “neuronal” or “glial”, respectively. Edges connecting “neuronal” components with “glial” components were discarded.

### Tissue appearance normalization in phase2

The EM images in the phase2 region of the dataset were acquired with different microscope settings following an extended period of wafer storage, resulting in different SNR characteristics and visibly different tissue appearance, even after applying contrast normalization (CLAHE) Initial attempts to segment this part of the volume resulted in severe merge errors. We therefore decided to preprocess the images using SECGAN ^77^, a machine learning technique designed to match the appearance (but not the semantic content) of 3d images from one source to another, so that impact on instance segmentation is minimized.

We trained three SECGAN models to make the phase2 part of the volume resemble phase1. The models processed data at 32×32×30 nm^3, 16×16×30 nm^3 and 8×8×30 nm^3 voxel size. Each SECGAN model used two independent 3d ResNet18 discriminators (one processing image data, and one processing segment probability maps produced by the FFN) and a residual convstack generator, with a depth of 8 modules and using only VALID convolutions. The models operating at 8×8×30 nm^3 and 16×16×30 nm^3 resolution were trained to map between a 32×32×60 µm^3 subvolume of phase1 and a 32×32×21 µm^3 subvolume of phase2. The model operating at 32×32×30 nm^3 resolution used a different pair of subvolumes (64×64×60 µm^3 for phase1 and 64×64×30 µm^3 for phase2), selected from a soma-dense region since the low-resolution FFN model was used for soma segmentation only.

After training, we used the FFN models to compute test segmentations of a small 500×500×448 vx^3 subvolume preprocessed with different checkpoints of the SECGAN, compared them to manual annotations, and selected the checkpoint minimizing the number of mergers for full volume inference. The phase2 images transformed with the SECGANs and the selected checkpoints were then used instead of the original EM as the input to the FFN when performing instance segmentation as outlined before.

### CAVE

We deployed a web-based proofreading tool, based on CAVE ^31^ (connectome annotation versioning engine), that allows collaborative proofreading of the Fish 1.0 volume directly in the browser. CAVE allows the assembly of neurons, which are split into many pieces in the automated segmentation, and the correction of neuron objects that are falsely merged with other segments. Researchers worldwide can sign up to contribute to proofreading the latest segmentation version (see <u>website</u> for more details). New proofreaders can familiarize themselves with CAVE in a sandbox segmentation and, after a successful assessment, contribute to the production segmentation. We also provide a <u>programmatic interface</u> to CAVE that lets users create and view custom annotation tables in CAVE, download the latest mesh versions, and view the proofreading history through a changelog website.

### Synapse annotation and predictions

Directly annotating synapses in the target zebrafish EM volume is time consuming. Therefore we leveraged a semi-automatic approach to first predict synapses using a pretrained model and then proofread the automatic labels. Specifically, we trained a 3D U-Net ^78^ using the synapse labels in an EM volume of adult rat ^79^ as the base model. We further fine-tuned the model through a semi-supervised learning approach by combining the original labeled training volume with pseudo-labeled chunks (via the base model) from the zebrafish EM volume to improve the adaptation to the target volume. All the modeling scripts are implemented with the open-source PyTorch Connectomics ^80^ framework. When running inference on the zebrafish images, we lower the probability threshold to encourage more synapse predictions since the manual effort to remove a false positive prediction is significantly less than manually navigating in the dense 3D image volume and drawing a synapse mask from scratch. Our model generated 21,346 synapse predictions in seven regions of interest (ROI, 16.3×16.3×3.3 µm^3) of the target EM volume. After proofreading in VAST using a custom-written Matlab navigation script and subsequent mask correction, we curated 14,378 ground-truth labels, which is later used as training data to scale model-based synapse labeling to the whole volume.

The 14, 378 ground-truth labels were used to train an initial synapse model on the 8nm resolution EM data. The model was based on a 3D U-Net architecture, with 3 down stages, 3 up stages, an initial feature size of 32, and a scale factor of 2 per UNet stage. Each convolution kernel was 3×3×3 (XYZ), and maxpool stride was 3×3×1. A final fully-connected layer ended in a three-class classifier, where 0 was background, 1 was presynaptic, and 2 was postsynaptic after applying softmax. The model was trained at a batch size of 1, and augmentations of reflection and permutations were applied to XYZ and XY, respectively. Each training example was calculated as the geometric centroid of the label mask. In order to help generalization during inference where FoVs may not be centered directly on a synapse, an additional augmentation of a random offset of between zero and +/-216×216×90nm was added to each training example. Due to the overwhelming ratio of background voxels to positive labels (approximately 14:1), we introduced a loss scaling of 2x for positive voxels in order to bias the network towards better recall. Training was performed for 80M steps.

Predictions were manually inspected for any systematic errors, and three more bounding boxes were added to improve recall (32.8×32.8×3.3µm^3 ROI size), which resulted in additional 5112 synapse annotations for GT. These additional annotations were added to the 14k GT examples for a total of 20k human-generated/-proofread examples. Using the same parameters as the U-Net above, the final model was trained for 4.09M steps, with the difference in training steps due to potential overtraining of the first model.

Inference was performed on the full volume. We ran connected components to assign unique ids to all predicted supervoxels. Synaptic pairs were performed by a multi-step connectome assembly pipeline. First a filtering step applied the tissue type classification masking model to ensure synapses were only present on somas or neuropil. A subsequent filtering step removed any predicted sites that were 30 voxels or smaller. Any remaining sites that spanned two or more neuropil segments at a threshold of a minimum of 20% of the original synaptic site’s supervoxels overlapping a given neural segment were split into unique sites with the same pre-or post-class and assigned unique IDs. A final filtering stage that removed any sites with a volume below 100 voxels, ensuring that any splits that introduced small or spurious new supervoxels were dropped. Finally we performed pairwise assignment of synaptic pre-and postsynaptic sites into synapses, defining a pair where the minimum euclidean distance between sites was 100nm or lower. This step intentionally did not enforce a 1:1 pre-/post-synaptic site pairing to allow for identification of potential polyadic synapses.

On rare occasions the model correctly predicted the presence of a synapse but reversed the pre-and post-classes. To correct these instances, we applied a synapse reorientation process as described in ^2^, identifying 4,973,737 synapses that required flipping, for an inverted prediction occurrence rate of 10.7%

### E/I classification model

491 *vglut2a*-and 547 gad1b-positive neurons were inspected for the presence of an axon with no merge errors, yielding a ground truth set of over 69,000 synapses (31,013 from *vglut2a* and 38,312 from *gad1b*-positive neurons) In total the ground truth consisted of 38,312 inhibitory and 31,013 excitatory synapses split into 80/20 train/test examples, with training examples upsampled and balanced to 306,490 examples of each class. We trained a ResNet model based on resnet50 ^2^ with a final fully-connected layer resulting in two classes: excitatory or inhibitory. Model inputs were two channels, with the first being the 16nm-scale uint8 EM data converted to float via z-score with a mean of 128 and standard deviation of 33, and the second being a trinary mask of the segments associated with each synapse during synapse assembly (see previous section) where the background was 0, pre-synaptic segment mask weighted 0.95 and post-synaptic mask weighted - 0.95. Input size was a patch shape of 1600×1600×720nm^3. Augmentations were the same as the U-Net model used during synapse identification, with the exception of the random movement parameter modified to be between zero and +/-272×272×120nm. Training used a batch size of 16 for 22M steps. Evaluating on the 20% held out synapses the model correctly identified 11,571 of 13,866 examples, for an accuracy of 83.45%.

### Synaptic polarity index computation

To estimate the polarity bias of each presynaptic neuron, we computed a polarity index using a Bayesian model comparing three possibilities for the synapse type composition: excitatory (E), inhibitory (I), and non-polarized/other (O). Let *n_exc_* and *n_inh_*represent the number of excitatory and inhibitory synapses made by a neuron, respectively. The likelihoods under each possibility were computed as follows:

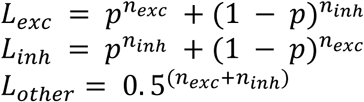

We used p=0.8 as the accuracy of the model given that the neuron is excitatory or inhibitory. The likelihoods are normalized to form posterior probabilities, assuming a uniform 1/3 prior for all three classes:

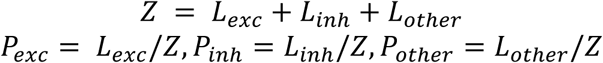

The final polarity index is computed as:

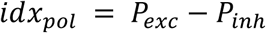

This index ranges from −1 (strongly inhibitory) to +1 (strongly excitatory), with values near 0 indicating mixed or ambiguous synaptic polarity. Synaptic input distributions across zebrafish brain regions To quantify the polarity of the synaptic input received by individual neurons, we identified all presynaptic inputs of each target cell and classified them as excitatory, inhibitory or other based on the computed polarity index (*idx_pol_*). Presynaptic cells and axon fragments with at least 4 synapses were included and were categorized as excitatory if *idx_pol_* ≥ 1/3, inhibitory if *idx_pol_* ≤ −1/3, and as “other” in the remaining cases. For each postsynaptic cell, we computed an input EI drive index defined as the normalized difference between excitatory and inhibitory incoming synapse counts. We also computed an input O drive index defined as the normalized difference between “other” and the summed excitatory and inhibitory incoming synapse counts. The index was aggregated by brain region based on MECE masks, and region-level statistics such as mean and standard error were computed. We computed these statistics with different sets of synapse number thresholds (**SFig. 6**), and compared to shuffled controls where the categories are randomly assigned to presynaptic cells and axon fragments (**SFig. 7**).

### Bootstrap-based propagation of polarity and orientation labels

To infer the orientation and polarity labels of unlabeled posterior lateral line neurons (PLLn) and medial octavolateralis nucleus (MON) neurons, we implemented a bootstrap-based label propagation algorithm that leverages partial ground truth and synaptic connectivity data. All analyses were performed separately for each brain hemisphere.

For both orientation and polarity inference, ground truth-labeled PLLn were separated into two groups based on molecular identity: vertical vs horizontal, and positive and negative. At the start of each bootstrap iteration, an equal number of seeds from each group were randomly sampled (by subsampling the larger group to the size of the smaller), ensuring balanced initial conditions and unbiased label propagation. This balancing procedure was reapplied after each label propagation cycle.

A synaptic connectivity matrix was constructed, with rows and columns corresponding to all included neurons and entries indicating synapse counts. To assess the robustness of the results given the estimated false-positive rate in synapse prediction, 15% of synapses were randomly removed at the start of each bootstrap iteration. This controlled perturbation introduced noise while testing the stability of the algorithm under realistic error conditions.

Each bootstrap iteration repeats the following label propagation cycles until convergence: Forward propagation: for every MON, a normalized index was calculated to quantify the relative strength of synaptic input from the two seed groups. Below, *N_group_* denotes the total number of synapses received from each seed group

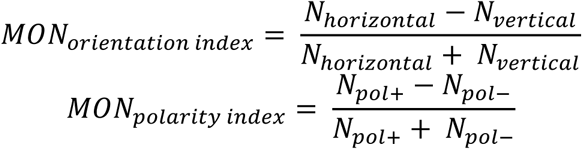

Backward propagation: for each PLLn, an *afferent score* was computed by summing its synaptic outputs to all MONs, weighted by each MON’s current index (orientation or polarity). This score estimates each PLLn’s affinity for orientation or polarity.

After each propagation cycle, PLLns with afferent scores above a positive threshold were provisionally assigned to one group, and those below a negative threshold to the other. The numbers in each group were again balanced by random subsampling. These forward and backward steps were repeated, updating group assignments each time, until convergence was achieved— defined as a change of less than 10^−6^in the binary assignment vector between consecutive iterations. This strict criterion ensured that further iterations would not meaningfully alter assignments. Upon convergence, the final assignments were used to recompute MON indices and PLLn afferent scores using the fully labeled set.

The entire propagation and label assignment procedure was repeated for 100 bootstrap samples. For each neuron, the fraction of bootstrap runs in which it was assigned to a given group was used as a probabilistic estimate of its orientation or polarity label. This provided uncertainty quantification for every assignment, reflecting both annotation noise and the structure of the underlying connectivity. Notably, the algorithm allows label updates when connectivity patterns conflict with seed assignments, but such reversals were rare: none of the 24 polarity-labeled seeds changed, and only 2 out of 52 orientation-labeled seeds were reassigned.

---

**SFig. 1.**
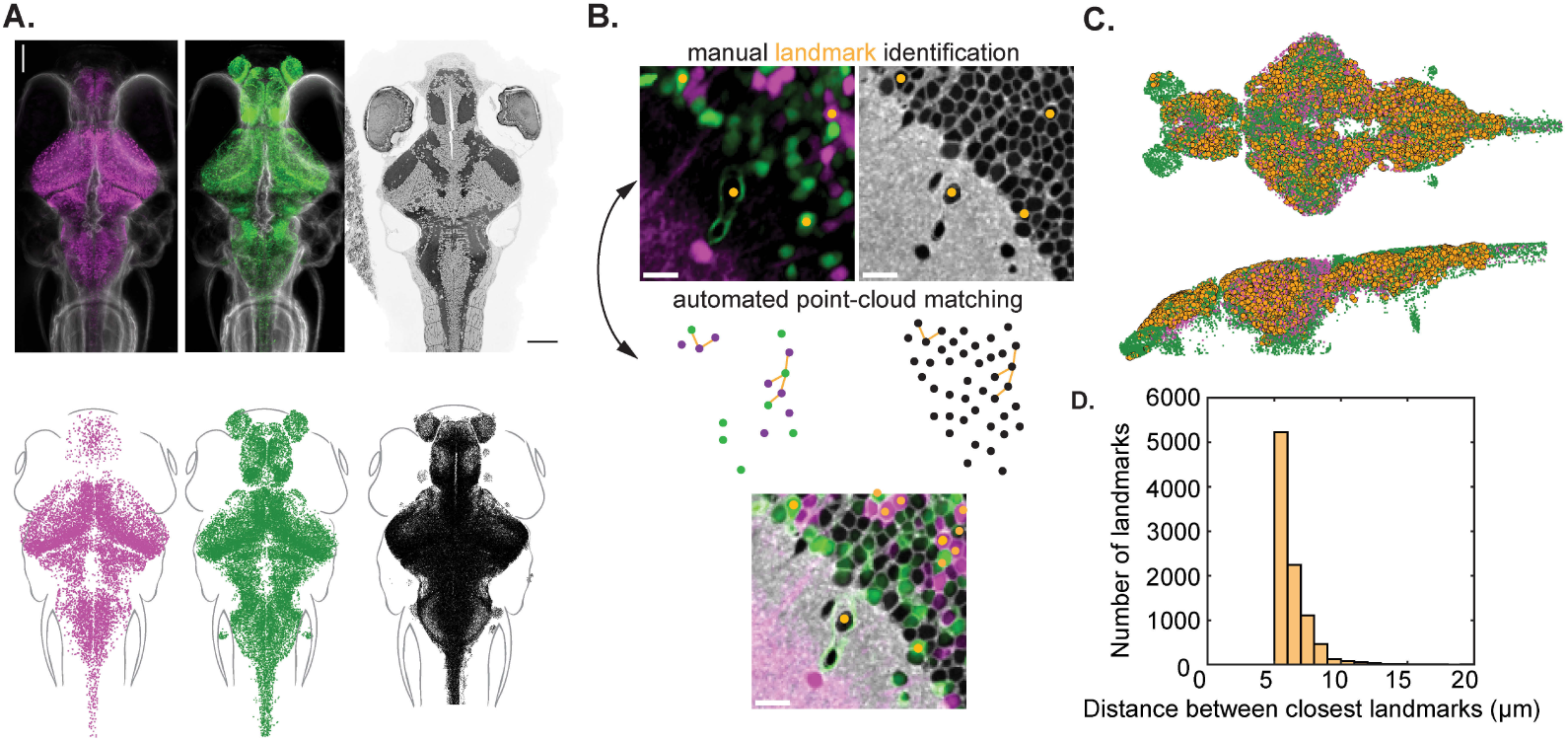
Correlated Light and Electron Microscopy (CLEM) approach. A, Maximum intensity projections of the *gad1b* and *vglut2a* confocal volumes (left) alongside a single electron microscopy (EM) section (scale bar = 100 µm). The image volumes are segmented into point clouds consisting of 10,937 *gad1b*, 24,812 *vglut2a*, and 187,053 EM cells. **B,** Semi-automated iterative process for identifying corresponding landmarks between the light microscopy (LM) and EM images. Initial manual landmarks are used to align the point cloud datasets. Groups of three or four LM cells are matched to EM cells in their local neighborhood, allowing for small rotations and translations of the group. Matches are added to the landmark point list, and the process is repeated with the updated landmark list. Manual landmarks can be added at any iteration. **C,** Distribution of the final set of 9,421 LM landmark cells used to transform the volumes. This final set is downsampled to a spacing of 10 pixels (5 µm) between landmarks for computational efficiency. **D,** Distribution of the distances between the two closest pairs of landmarks. The typical cell diameter is approximately 5 µm.

**SFig. 2.**
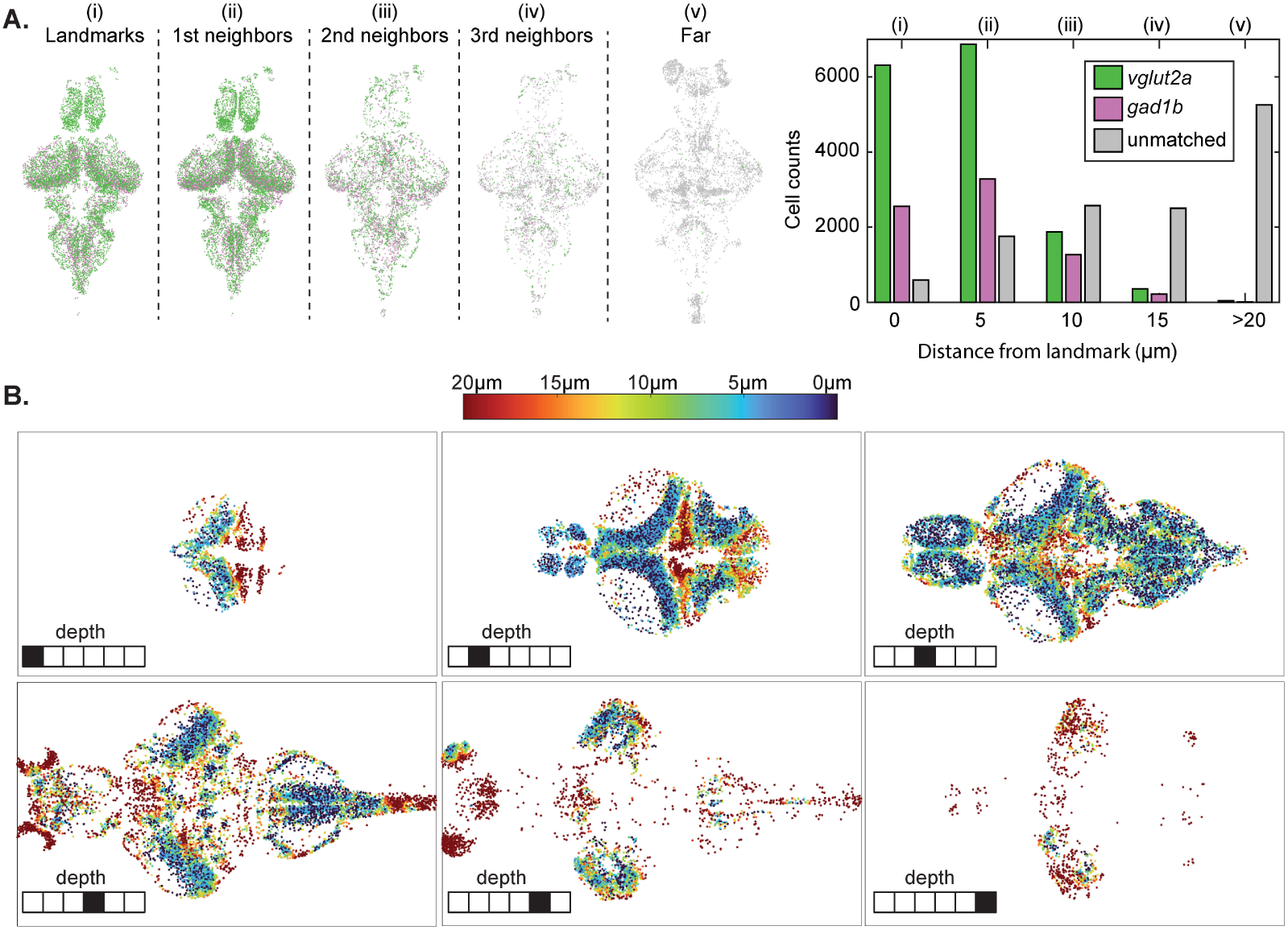
Correlated Light and Electron Microscopy (CLEM) quality. A, The registration quality for all 35,749 LM cells is evaluated after mapping the LM volume onto the EM volume using the 9,421 landmarks. Left: The cells are split into groups based on the distance from their centroid to landmark cell centroids. The landmarks group is at a distance equal to zero. Right: Cells are assigned as matched (*vglut2a* or *gad1b)* based on a visual inspection of a 20 µm x 20 µm region surrounding the cell center, and unassigned if the visual inspection determines that the region cannot be unambiguously matched. Roman numerals mark the corresponding cell groups between the cell spatial visualizations on the left and the match quality assignment on the right. **B,** Spatial distribution of all 35,749 LM cells across six depths of the LM volume, with cells color coded based on their distance to a landmark.

**SFig. 3.**
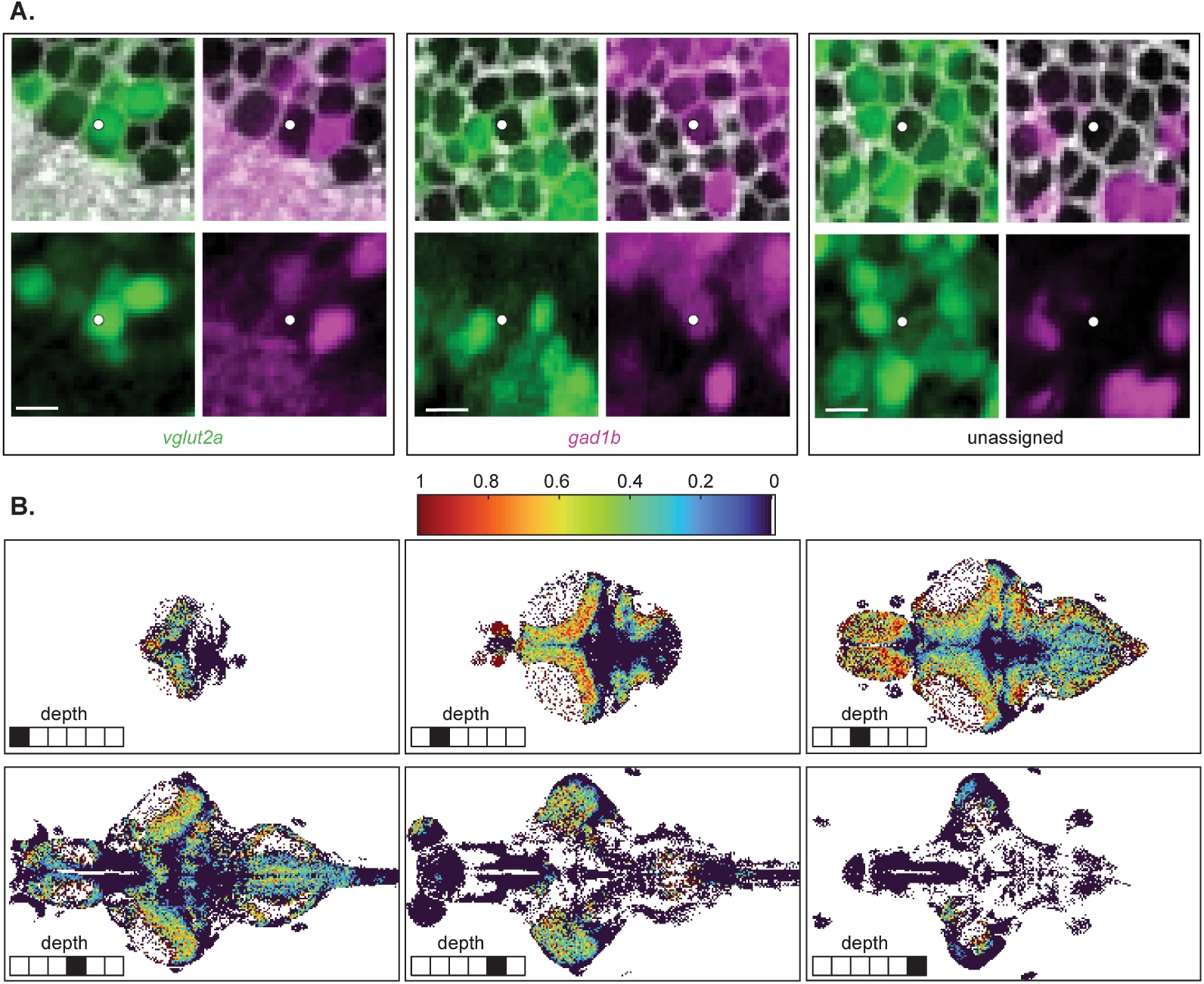
Intensity-based label assignment. A, Three example assignments based on 20 µm x 20 µm image overlays between the registered EM volume (grayscale) and the *vglut2a* (green) and *gad1b* (magenta) LM channels (scale bar = 5 µm). **B,** Spatial distribution of labeled cells across six depths of the EM volume, taken over 5 µm x 5 µm columns. White bar in the color scale indicates that background pixels are labeled in white.

**SFig. 4.**
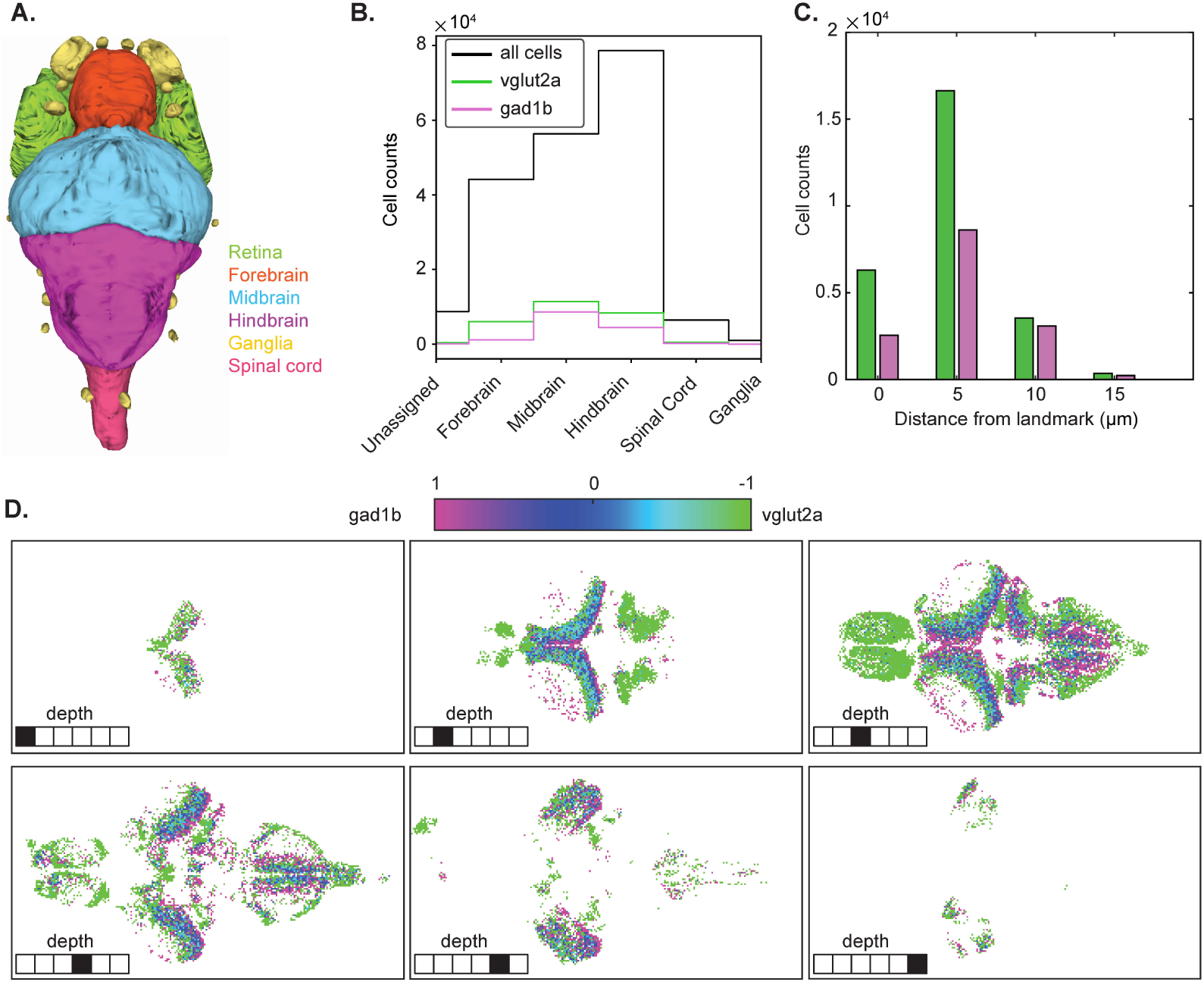
Excitatory and inhibitory neuron distribution after intensity-based manual label assignment. A, Major brain regions in the zebrafish, which are mutually exclusive and collectively exhaustive (MECE) as defined by the z-brain atlas (Vohra et al, Companion paper). **B,** Distribution of all labeled and registered *vglut2a* neurons (green), and *gad1b* neurons (magenta) across the major brain regions. **C,** Distribution of labeled cells *vglut2a* (green) and *gad1b* (magenta) according to distance from a landmark. **D,** Spatial distribution of label index across six depths of the EM volume, taken over 5 µm x 5 µm columns.

**SFig. 5.**
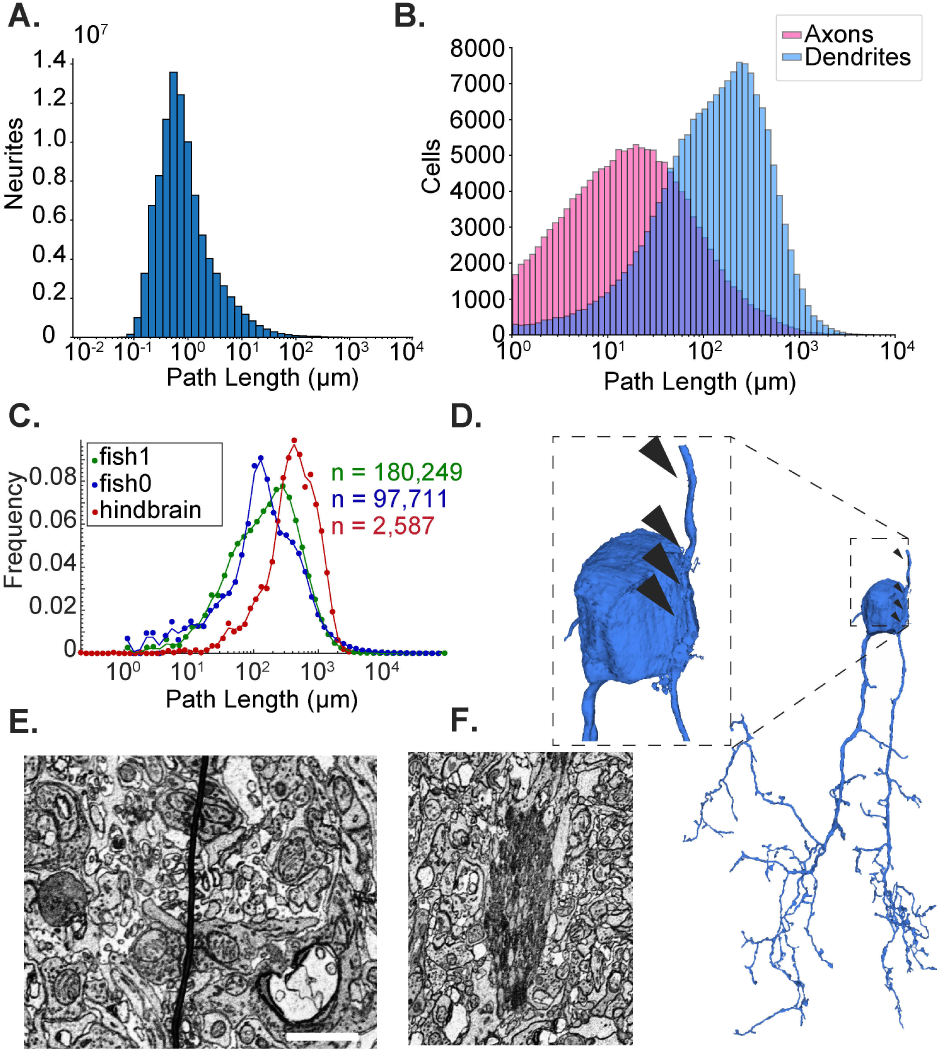
Quantification of automated segmentation and synapse quality. A, Distribution of neurite path lengths across the entire dataset; total reconstructed length sums to ∼250 meters. **B,** Distribution of axonal (pink) and dendritic (blue) neurite lengths assigned to somas. **C,** Comparison of neurite path length distributions from automated segmentations in this 7 dpf zebrafish brain (fish1, green), a previously published 5 dpf zebrafish brain (fish0, blue; ^9^), and a set of manually proofread neurons in the hindbrain of a zebrafish (red; ^30^). **D,** Example merge error where a passing neurite is erroneously joined to a soma (arrowheads). **E,** Example of tissue wrinkle; scale bar, 1 µm. **F,** Example of a staining artifact—axon bundle overstained with osmium; scale matched to **E**.

**SFig. 6.**
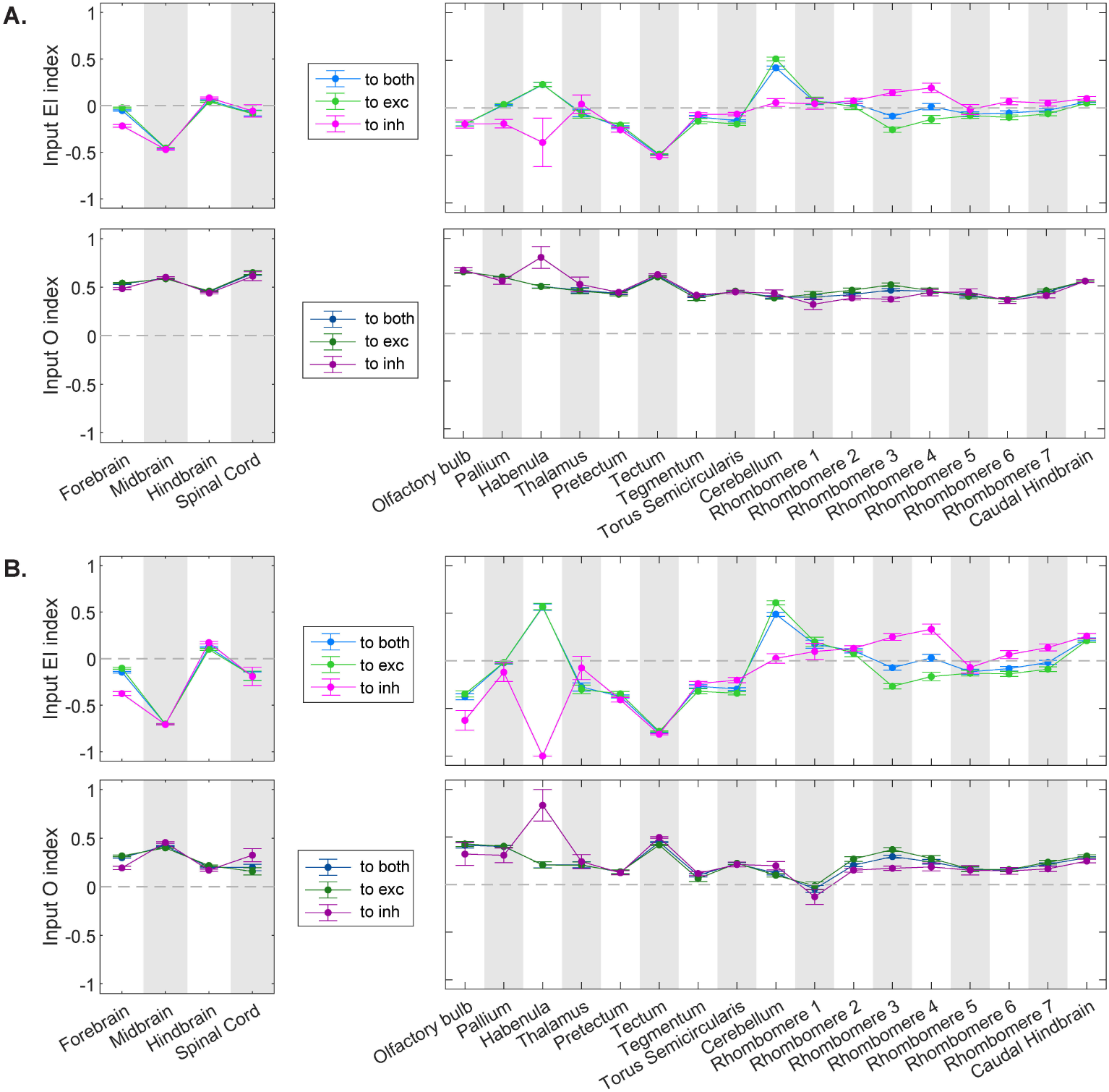
Threshold choice effect on the input drive index. Average input drive indices to the 41,000 *vglut2a* and *gad1b* neurons segregated by brain region computed with different synapse thresholds: **A.** no threshold, **B.** at least 10 synapses. Error bars reflect standard error of the mean.

**SFig. 7.**
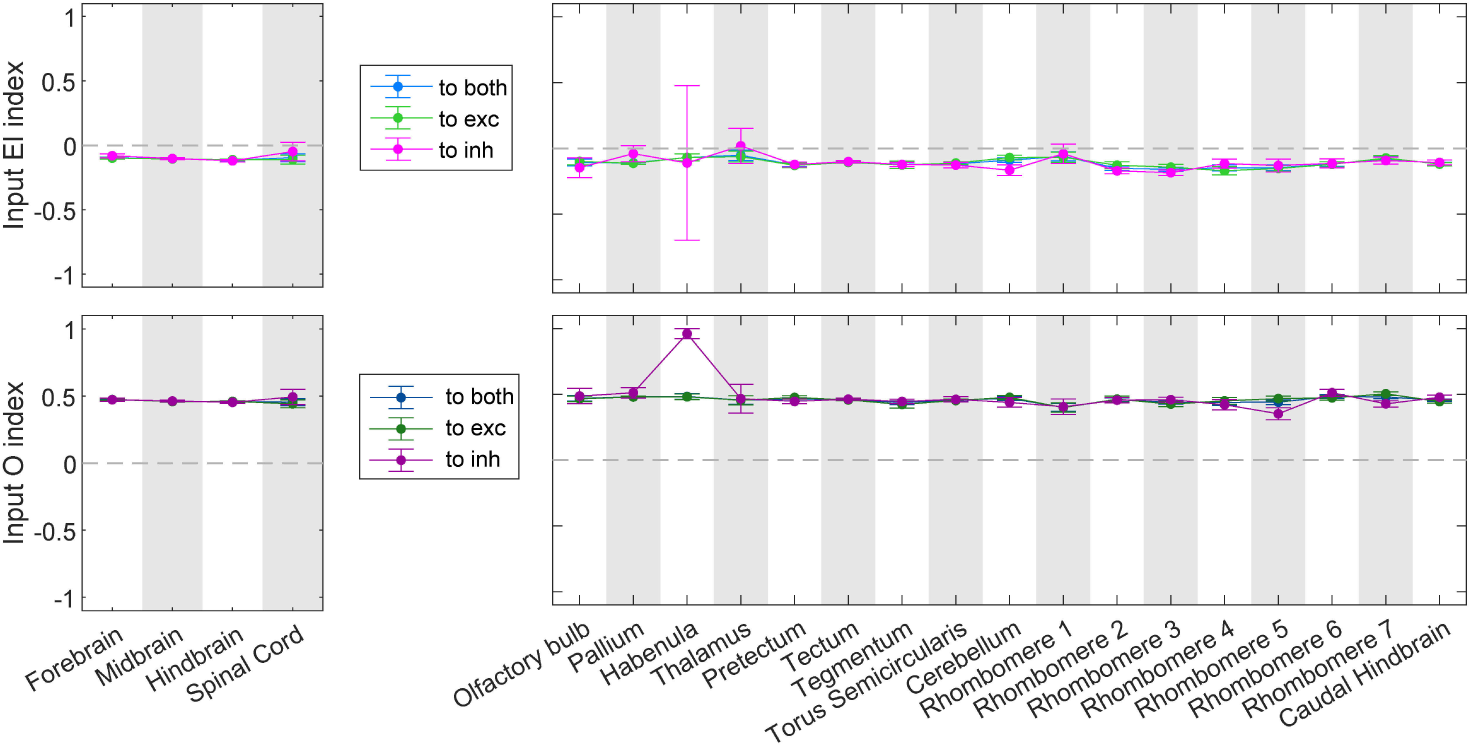
Shuffled control effect on the input drive index. Average input drive indices computed with shuffled polarity indices for the input synapses to the 41,000 *vglut2a* and *gad1b* neurons segregated by brain region (threshold is at least four synapses). Error bars show standard error of the mean.

**SFig. 8.**
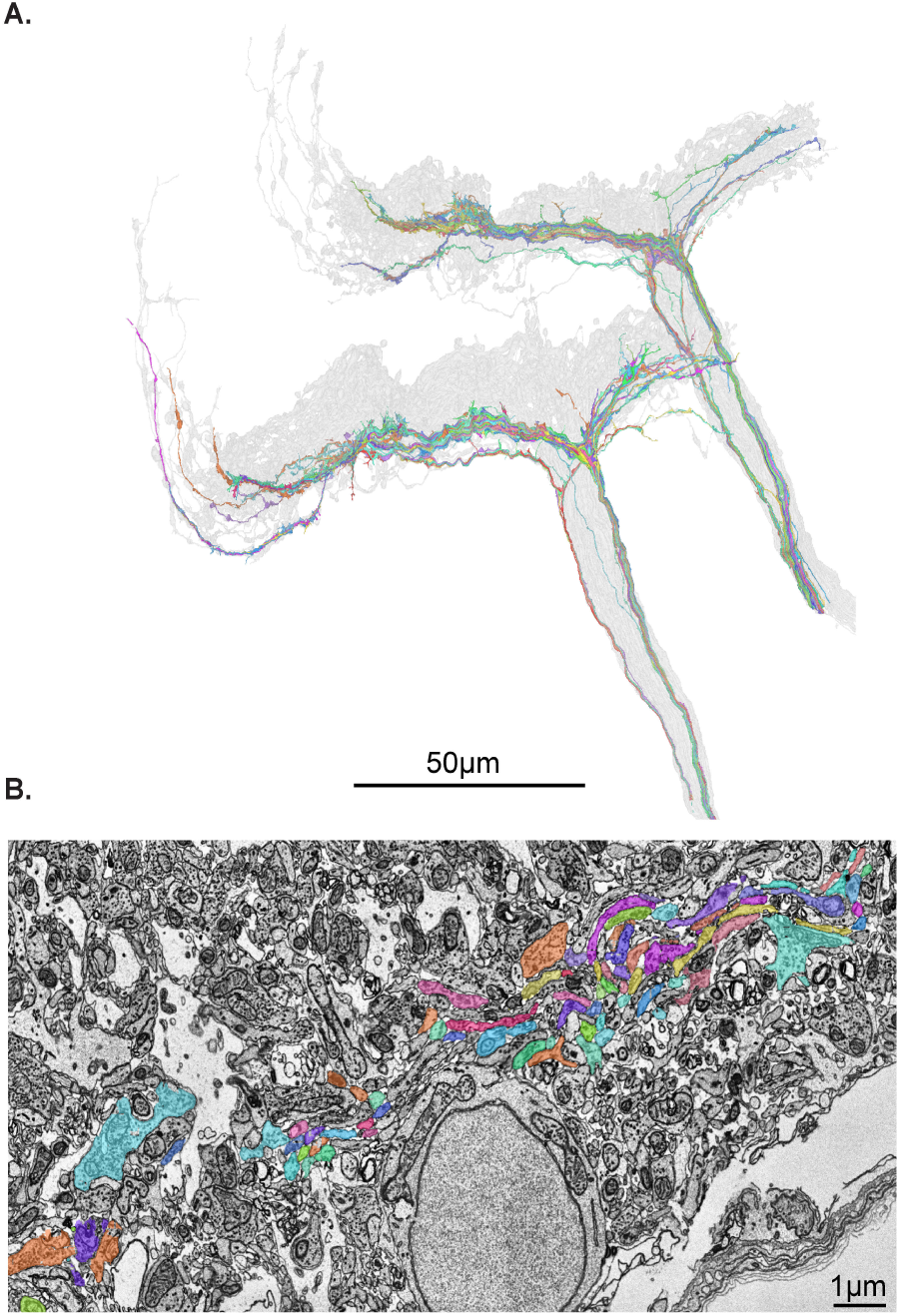
Sheet-type afferent neurons. A, Reconstructions of 152 bulb-type afferent arborizations (gray) and 102 sheet-type afferent arborizations (colors) **B,** Ultrastructure of the sheet-type afferens, which travel in bundles and have few synapses.

